# Angiotensin 1-7 Modulates the Dynamics and Activation of the Proto-oncogene Mas Receptor

**DOI:** 10.1101/2025.07.07.663239

**Authors:** Ekrem Yasar, Segun Dogru, Erol Eroglu, Nazmi Yaras

## Abstract

The proto-oncogene Mas receptor (MasR, UniProt ID: P04201) is a class-A (orphan-type) G protein-coupled receptor (GPCR) that mediates the protective effects of Angiotensin 1-7 (Ang 1-7) within the renin–angiotensin system (RAS). Despite its therapeutic relevance, the molecular mechanisms underlying MasR activation by Ang 1-7 remain elusive due to the lack of experimental structural data. In this study, we performed 1-microsecond all-atom molecular dynamics (MD) simulations of AlphaFold-modeled active and inactive MasR conformations, with and without Ang 1-7, to characterize ligand-induced conformational dynamics.

Ang 1-7 binding led to increased interaction stability in the active state, reflected by higher occupancy of hydrogen bonds, salt bridges, and hydrophobic contacts. Structural analyses revealed reduced RMSD/RMSF values and stabilization of key transmembrane (TM) helices and the NPxxY micro-switch. TM distance and dihedral analyses indicated partial TM6 displacement and time-dependent NPxxY reorganization. Network-based metrics including betweenness centrality and shortest path length highlighted the emergence of state-specific communication hubs, while PCA, correlation and communication propensity analyses revealed enhanced conformational diversity and selective inter-residue signaling in the ligand-bound state. Molecular mechanics Poisson–Boltzmann surface area (MM/PBSA) calculations showed favorable binding energetics in the active state (ΔG_Bind_ = –13.99 kcal/mol).

These results demonstrate that Ang 1-7 acts as a partial agonist of MasR by stabilizing the inactive conformation while inducing limited activation features in the active state through non-canonical micro-switch dynamics. This work advances structural insights into MasR regulation and provides a foundation for therapeutic targeting of the ACE2/Ang 1-7/MasR axis.

## 1 INTRODUCTION

The renin angiotensin system (RAS) is a central hormonal pathway that regulates blood pressure, fluid balance, and vascular tone, playing a crucial role in cardiovascular and renal homeostasis (1,2). The classical axis of RAS, comprising Angiotensin Converting Enzyme (ACE), Angiotensin II (Ang II), and the Angiotensin II type 1 receptor (AT1R) (ACE/Ang II/AT1R), is well established for its vasoconstrictive, pro-inflammatory, and pro-fibrotic effects (3,4). In contrast, a counter-regulatory pathway known as the protective arm of the RAS has gained prominence. This axis involves ACE2, Angiotensin 1-7 (Ang 1-7), and the Mas receptor (MasR) (ACE2/Ang1-7/MasR), and is associated with vasodilation, anti-inflammatory signaling, anti-fibrotic actions, and cardioprotective outcomes (5,6).

Ang 1-7, a heptapeptide (Asp-Arg-Val-Tyr-Ile-His-Pro, DRVYIHP) primarily generated by ACE2 through cleavage of Ang II, functions as the main ligand of MasR, an orphan-type G-protein coupled receptor (GPCR) encoded by the proto-oncogene MAS1 (7). MasR is predominantly expressed in the heart, kidneys, vasculature, lungs, and several endocrine tissues, where it contributes to endothelial nitric oxide (NO) production, inhibition of oxidative stress, and anti-proliferative signaling (8). Upon Ang 1-7 binding, MasR activation leads to phosphorylation of endothelial nitric oxide synthase (eNOS), reduced nikotinamid adenin dinükleotid fosfat (NADPH) oxidase activity, and downregulation of Nuclear Factor kappa B (NF-κB) and mitogen-activated protein kinase (MAPK) pathways, which collectively protect against vascular damage and remodeling (9).

Numerous experimental studies have highlighted the physiological and therapeutic relevance of the ACE2/Ang 1-7/MasR axis. In preclinical models of myocardial infarction and heart failure, Ang 1-7 administration preserved ejection fraction, reduced cardiac hypertrophy, and prevented pathological remodeling by activating PI3K/Akt and AMP-activated protein kinase (AMPK) pathways (10). MasR-deficient mice failed to exhibit these effects, confirming the receptor’s essential role in transducing Ang 1-7’s cardioprotective actions (11). In hypertensive rats, Ang 1-7 improved endothelial function and attenuated vascular hypertrophy by enhancing NO bioavailability and suppressing pro-oxidant signaling (12). Additionally, Ang 1-7 treatment inhibited fibrosis in cardiac and renal tissues through suppression of TGF-β1 and collagen deposition, reinforcing its role in tissue remodeling and homeostasis (13).

Beyond the cardiovascular system, the MasR–Ang 1-7 axis has demonstrated anti-tumor potential in a growing number of studies. MasR expression has been detected in several malignancies, including breast, prostate, and lung cancers, as well as glioblastoma (14,15). In experimental cancer models, Ang 1-7 reduced tumor volume, microvascular density, and cell proliferation by inhibiting vascular endothelial growth factor (VEGF), matrix metalloproteinase-9 (MMP-9), and ERK1/2 phosphorylation (16). Moreover, the axis has been implicated in attenuating epithelial-to-mesenchymal transition (EMT) and enhancing chemosensitivity, underscoring its promise as a therapeutic target in oncology (17).

Despite these functional insights, the molecular mechanisms underlying MasR activation remain poorly defined. Unlike well-studied class-A GPCRs such as β2-adrenergic and AT1 receptors, MasR lacks an experimentally resolved three-dimensional structure. This has hindered efforts to elucidate its conformational transitions, ligand recognition mechanisms, and signal initiation pathways. Moreover, contradictory findings regarding whether Ang 1-7 binds MasR directly or through accessory partners further complicate mechanistic interpretation (18,19).

In this context, computational modeling serves as a critical tool to bridge structural gaps and uncover receptor activation mechanisms. Recent advances in structure prediction tools, particularly AlphaFold (20), have enabled the generation of high-confidence models for both active and inactive conformations of proteins such as GPCRs, including those lacking experimental structures. When integrated with molecular dynamics (MD) simulations, these models allow for the exploration of receptor flexibility, ligand-induced conformational changes, and structural transitions associated with activation. In class-A GPCRs, activation is commonly associated with highly conserved micro-switch rearrangements, such as the outward movement of TM6, the reorganization of the NPxxY motif on TM7, and the breaking of the ionic lock between TM3 and TM6 features that underpin the switch from inactive to active states and facilitate G-protein coupling (21). For orphan GPCRs like MasR, whose endogenous activation mechanism remains elusive, such modeling efforts are particularly valuable (22). Leveraging AlphaFold-predicted conformers, MD simulations offer a powerful framework to dissect ligand–receptor interactions, assess structural plasticity, and gain mechanistic insights into activation processes, thus advancing the molecular characterization of poorly understood receptors such as MasR.

Nevertheless, computational investigations of MasR remain limited. Prokop et al. modeled MasR using I-TASSER and performed short-time MD simulations with docked Ang peptides, focusing only on binding pose stability (23,24). A more recent study by Yasar et al. employed 300-ns MD simulations to compare MasR in the presence and absence of Ang 1-7, identifying preliminary dynamic differences (25). However, neither study explored activation-related conformational states, nor did they examine long-timescale dynamics necessary to resolve microswitch transitions or state-specific modulation.

To address these limitations, we conducted an in-depth computational study to investigate how Ang 1-7 binding influences the conformational dynamics and signal transduction mechanisms of the MasR. Using MD simulations based on AlphaFold-predicted active and inactive MasR structures, we assessed the effects of Ang 1-7 on receptor stability, residue-level communication, and structural rearrangements. Key analyses included global and local flexibility assessments, interaction network mapping, and principal motion decomposition, enabling the identification of dynamic communication pathways and functionally relevant residues. These integrative approaches provided a coherent framework for characterizing Ang 1-7 induced modulation of MasR across different receptor states. Our findings provide mechanistic insight into how Ang 1-7 modulates MasR activity and offer a valuable foundation for future studies exploring the therapeutic relevance of this axis in cardiovascular and oncological contexts.

## 2. MATERIALS AND METHODS

### 2.1. Structural Preparation

To investigate the structural basis of Ang1-7 modulation of the human Mas receptor (MasR, UniProt ID: P04201), active and inactive conformations were retrieved from GPCRdb (26), predicted by AlphaFold with high confidence (pLDDT >90), requiring no structural completion (20). The absence of an experimentally resolved structure in the Protein Data Bank (PDB) (27) necessitated the use of these computational models. The NMR structure of Angiotensin 1-7 (Ang1-7) was obtained from PDB (ID: 2JP8) (28) and fallowing by protonation states assignment at pH 7.0 using CHARMM-GUI (29) for ligand and receptor. Blind peptide-receptor docking was performed using MDockPeP (30), a statistical potential-based ab initio docking platform, with ITScorePeP scoring function. For each conformation, the top-scoring pose (lowest ITScorePeP) was selected as the initial structure for MD simulations (Fig 1).

**Figure 1.**
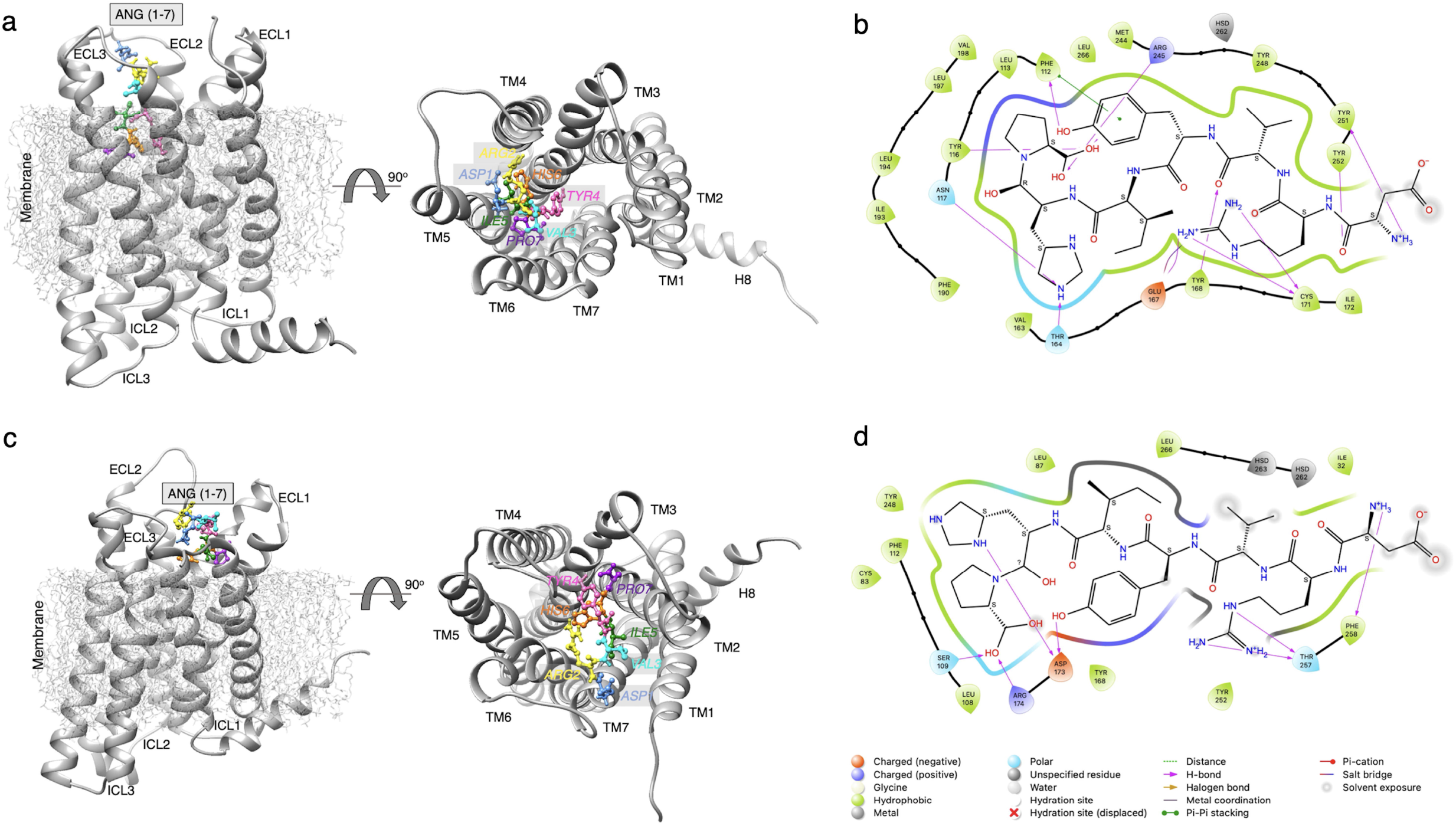
Binding orientation and interaction analysis of Ang 1-7 with MasR in inactive and active conformations. a) Side and top views of Ang 1–7 docked into the inactive-state MasR structure modeled via AlphaFold and obtained from GPCRdb. b) 2D interaction diagram showing key residues involved in ligand binding in the inactive state. c) Side and top views of Ang 1–7 docked into the active-state MasR structure. d) 2D interaction map highlighting the distinct interaction network in the active conformation.

### 2.2. Molecular Dynamics Simulation Setup

Four systems were constructed: Apo-MasR (active), Apo-MasR (inactive), Ang1-7-MasR (active), and Ang1-7-MasR (inactive). Each receptor or receptor-ligand complex was embedded into a symmetric lipid bilayer (POPC:POPE:CHL, 2:2:1) using CHARMM-GUI Membrane Builder (31), solvated with ∼30,000 TIP3P water molecules (32) in a 9.5 nm × 9.5 nm × 10 nm box, and neutralized with 0.15 M NaCl (∼100,000 atoms total). Simulations utilized the CHARMM36m force field (33) in GROMACS 2020.5 (34), executed on NVIDIA RTX A4000 GPUs. Systems underwent energy minimization via steepest descent and a six-step equilibration protocol (1 ns NVT at 310 K with 10 kcal/mol·Å^2^ restraints, 2 ns NPT with gradual restraint release), followed by 1-µs production runs with a 2-fs time step. Temperature was maintained at 310 K using the Nose-Hoover thermostat (35), pressure at 1 atm with the Parrinello-Rahman barostat (36), and periodic boundary conditions, Particle Mesh Ewald electrostatics (37), and LINCS bond constraints (38) were applied.

### 2.3. Structural and Interaction Based Analyses

Trajectory analyses assessed ligand-induced and receptor-intrinsic dynamics across all four systems. Root-mean-square deviation (RMSD) and fluctuation (RMSF) were calculated using GROMACS tools (39), with RMSD computed for the full protein, individual TM helices, and Ang1-7 over 10000 frames. Two-dimensional ligand-receptor interaction maps were generated using Schrödinger Maestro (2020 demo version) (40). Hydrogen bond occupancies were determined over 10000 frames using VMD 1.9.3 (41), with a 3.5 Å distance and 30° angle cutoff. Salt bridge interactions were calculated over 2000 frames with a 4 Å distance cutoff, and hydrophobic interactions (4 Å cutoff) with frame-wise occupancy were computed using custom Python scripts (https://github.com/eygpcr/hydrophobic_interactions) based on MDAnalysis (42). In addition to all trajectory files, movies of ligand-receptor interactions, created in VMD, are deposited on Zenodo (https://doi.org/10.5281/zenodo.15781385). Center-of-mass (COM) distances between intracellular ends of TM3, TM5, TM6, and TM7 were measured using the gmx distance module to evaluate hydrophobic lock rearrangements. Dihedral angles (φ, ψ, χ_1_) dynamics of the NPxxY motif were analyzed using in-house Python tools (https://github.com/eygpcr/microswitches_dihedral_angles) based on MDAnalysis, focusing on N276, P277, F278, I279, and Y280.

### 2.4. Binding Free Energy Calculations

Molecular Mechanics/Poisson–Boltzmann Surface Area (MM/PBSA) is a widely used method for estimating the binding free energy of protein–ligand complexes (43). In this study, full molecular dynamics trajectories (1 μs, 2000 frames per system) were used to evaluate the binding affinities of Ang 1–7 to both inactive and active conformations of the MasR.

Binding energy components were calculated using the gmx_MMPBSA package (44), which implements the following equation:

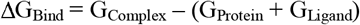

where G_Complex_ represents the total free energy of the protein–ligand complex, and G_Protein_ and G_Ligand_ denote the total free energies of the receptor and the peptide, respectively. Energy decomposition included van der Waals, electrostatic, polar solvation, and nonpolar solvation terms. A dielectric constant of 80 and a grid spacing of 0.5 Å were applied consistently across all systems.

### 2.5. Advanced Conformational and Signaling Network Analysis

Advanced analyses were conducted using the MDM-TASK suite (45,46) to identify key residues in conformational transitions, structural communication and signaling. Principal Component Analysis (PCA), Dynamic Cross-Correlation (DCC), Communication Propensity (CP), and Dynamic Residue Interaction Network (DRIN) (see Supplementary Methods for details) mapping employed ±3σ and ±2σ thresholds. Significant signaling residues were mapped onto 2D snake diagrams via GPCRdb. Per-residue fluctuation trends were visualized using VMD’s timeline plugin.

## 3 RESULTS and DISCUSSION

### 3.1. Angiotensin 1-7 Binding Interactions and Stability Profiles

Our 1-µs molecular dynamics (MD) simulations provided detailed insights into the binding interactions of Ang 1-7 with the MasR, an orphan type GPCR pivotal to the RAS protective-arm (47,48). The docking poses and two-dimensional (2D) interaction maps (Fig1, FigS2) revealed distinct binding modes, with Ang 1-7 adopting a compact orientation in the inactive state, engaging extracellular loops (ECL2, ECL3) and transmembrane (TM) helices (TM4, TM5, TM7), and a more outward configuration in the active state. These structural differences are driven by specific intermolecular contacts, quantified over 10,000 frames.

Hydrogen bond (H-bond) stability, assessed in Fig2a-b and detailed in TableS1, showed that Ang 1-7 forms more persistent interactions in the active state, with occupancies reaching 52.23% (SER109^3.29^-PRO7) (49) compared to a maximum of 35.45% (ARG245^6.52^-PRO7) in the inactive state, consistent with enhanced ligand-receptor affinity. Salt bridge persistence (Fig2c-d, TableS2) further supported this trend, with active-state occupancies of ∼10% (ASP1-GLU175) and ∼5% (ARG2-GLU175), compared to ∼6% (ASP1-ASP173) and ∼10% (ASP1-GLU175) in the inactive state, indicating a shift toward optimized electrostatic stabilization [34]. Hydrophobic interactions, depicted in Fig2e-f, were significantly stronger in the active conformation, with PHE112^3.32^-PRO7 reaching 84% occupancy (versus 46% inactive), alongside TYR248^6.55^-ILE5 and TYR95-ARG2 both at 55%, compared to 53% (ARG245^6.52^-PRO7) in the inactive state (Fig2f). These findings suggest that Ang 1-7’s binding affinity and stability are markedly enhanced in the active state, likely due to a better fit within the reorganized binding pocket, a common feature of GPCR activation (21,50).

**Figure 2.**
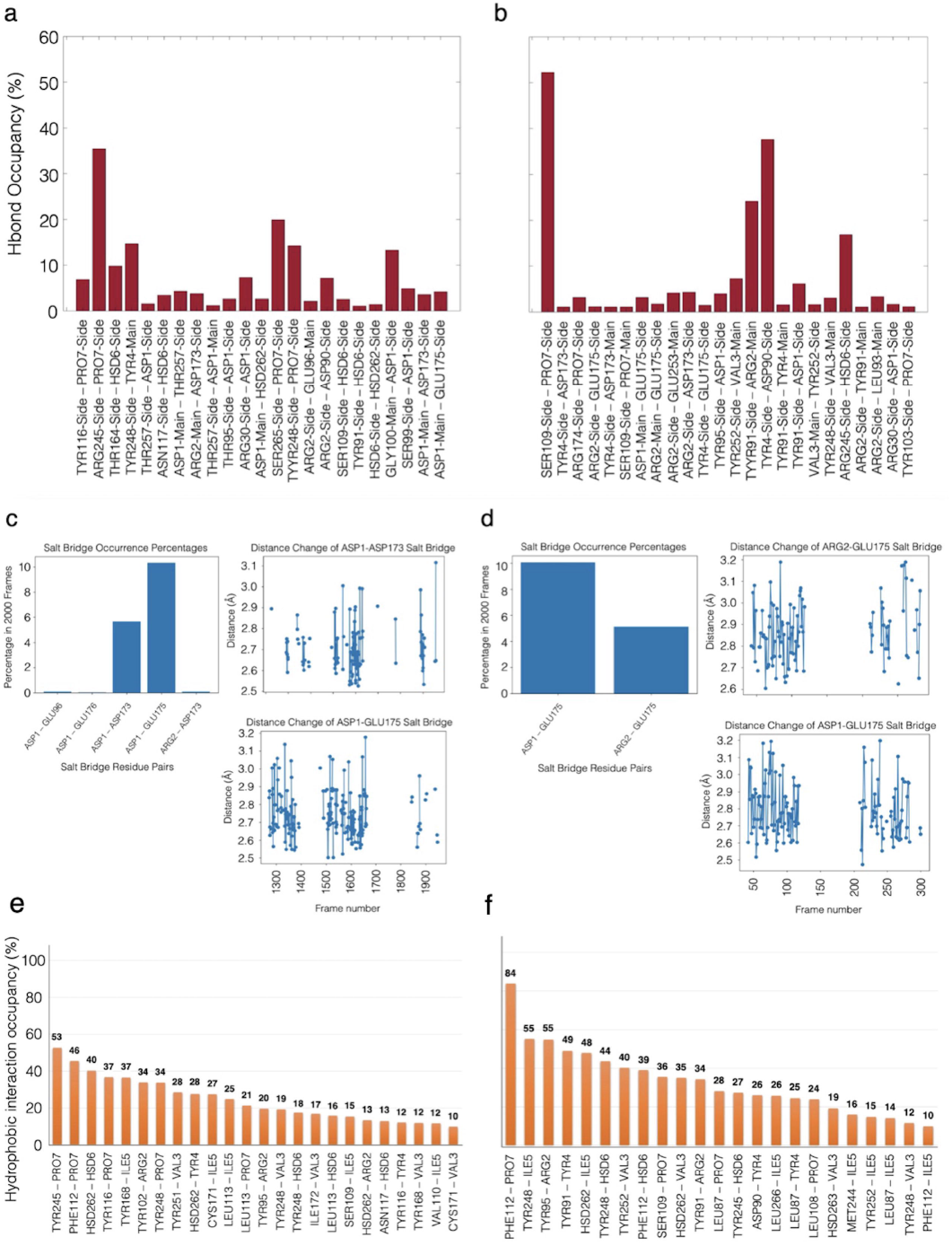
Intermolecular interaction analysis between Ang 1–7 and MasR in inactive and active conformations during MD simulations. a–b) Hydrogen bond occupancy between Ang 1–7 and MasR residues over the course of the simulations. Bar plots represent the percentage occupancy of each hydrogen bond in a) the inactive-state MasR and b) the active-state MasR. c–d) Salt bridge analysis between Ang 1–7 and MasR. Bar plots show the occurrence percentages of prominent salt bridges in c) inactive and d) active conformations. Line graphs display distance fluctuations of the most persistent salt bridges across selected trajectory frames. e–f) Hydrophobic interaction occupancy between Ang 1–7 and MasR residues. Bar plots illustrate the percentage of hydrophobic contact occupancy during the simulations for e) inactive and f) active MasR conformations.

The dynamic yet stable interactions observed, characterized by enhanced hydrogen bond, salt bridge, and hydrophobic occupancies, could be attributed to a modulated signaling profile that differs from the full activation typically associated with classical GPCR agonists (51-53). Studies suggest that Ang 1-7 serves as the primary endogenous ligand for MasR, exhibiting agonist properties such as vasodilation and anti-fibrotic effects (54,55), yet its activation mechanism remains context-dependent, with evidence pointing toward partial or biased agonism. This is supported by findings that the enhanced H-bond, salt bridge, and hydrophobic occupancies in the active state (Fig2) align with the partial agonist profile reported by (56-58), who noted that Ang 1-7 selectively activates specific pathways (e.g., ERK1/2, PI3K/Akt) without fully inducing classical G protein signaling. This nuanced role in MasR modulation contrasts with earlier docking studies lacking temporal resolution (23), underscoring the value of long-timescale MD simulations in capturing the ligand-induced stabilization mechanisms that underpin these functional differences.

### 3.2. Structural Dynamics and Flexibility Modulation by Angiotensin 1-7

The structural behavior of MasR in response to Ang 1-7 binding was explored through RMSD and RMSF analyses (Fig3), alongside TM helix and Helix 8 (H8) RMSD profiles (FigS1) and two-dimensional timeline visualizations (FigS2). In the inactive state, RMSD profiles indicated apo fluctuations of 3-5 Å, peaking at ∼7 Å after 600 ns, suggesting a late conformational shift that Ang 1-7 binding mitigated by stabilizing the structure at 4 Å, with ligand RMSD consistently at 3-4 Å (Fig3a). This points to Ang 1-7’s ability to enhance structural compactness, a key feature for maintaining receptor integrity. In the active state, apo RMSD held steady at 3 Å until a 6-7 Å peak emerged after 600 ns, whereas Ang 1-7 binding shifted the baseline to ∼5 Å with a similar peak, while ligand RMSD remained stable at 3-4 Å (Fig3c), indicating partial stabilization amid heightened flexibility. RMSF analysis revealed nuanced regional effects. In the inactive state, apo displayed elevated flexibility with 6 Å in ECL1 (87-104 residues), 5 Å in ECL2 (173-185 residues), and 4 Å in ECL3 (246-263 residues), which Ang 1-7 reduced to 3 Å, maintained at 5 Å, and lowered to 2 Å, respectively; notably, TM6 remained unchanged, while ICL2 and TM7 showed increased flexibility (Fig3b). This suggests Ang 1-7 selectively stabilizes extracellular loops, preserving intracellular dynamism. In the active state, apo RMSF values were 3 Å (ECL1), 5 Å (ECL2), 4 Å (ECL3), and 6 Å (H8, 284-298 residues), shifting with Ang 1-7 to 4 Å (ECL1), 5 Å (ECL2), 2 Å (ECL3), and 5 Å (H8), alongside an RMSF change between 1-4 Å in TM5/ICL3 (186-224) and a decrease from 3 Å to 1 Å in TM6 (225-245) (Fig3d). This pattern highlights ligand-induced stabilization in TM6 and ECL3, balanced by enhanced flexibility in TM5/ICL3, likely supporting G-protein interactions.

**Figure 3.**
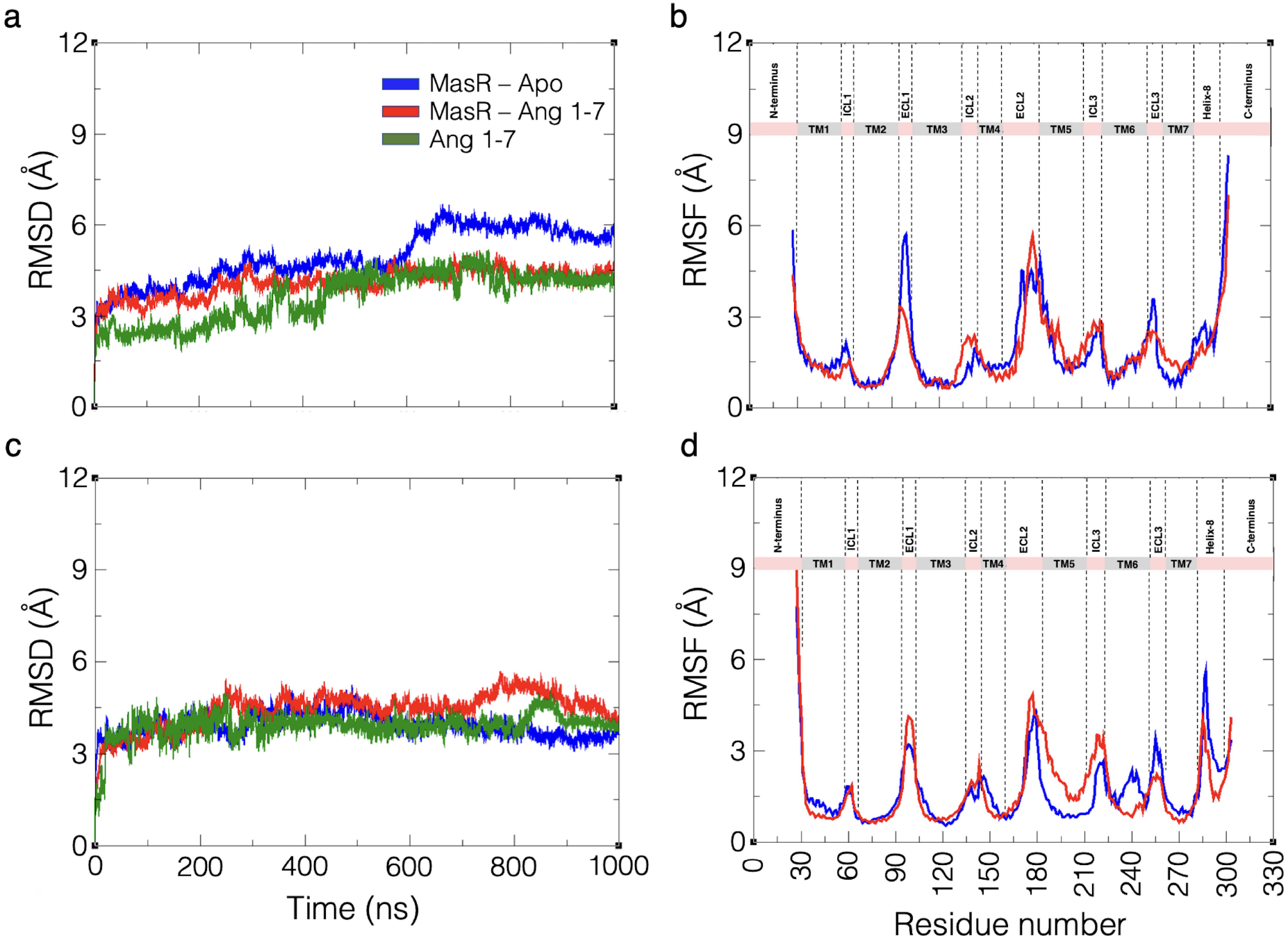
Structural dynamics of MasR in inactive and active conformations with and without Ang 1–7 binding. a) RMSD profiles of the inactive-state MasR over 1000 ns MD simulations. Blue: apo form; red: Ang 1–7-bound form; green: RMSD of Ang 1–7 ligand itself. b) RMSF analysis of inactive MasR, showing residue-level flexibility differences between apo (blue) and Ang 1–7-bound (red) states. c) RMSD profiles of the active-state MasR with the same color coding. d) RMSF comparison of the active-state MasR in apo and ligand-bound conditions.

These dynamics are further supported by timeline analysis, where apo states (FigS2a, c) exhibited heterogeneous color patterns reflecting high conformational variability, particularly in TM6, TM7, and H8, while Ang 1-7-bound states (FigS2b, d) showed more uniform distributions, indicating reduced heterogeneity. TM helix and H8 RMSD changes (FigS1) reinforced this, with Ang 1-7 attenuating fluctuations in TM6 and H8 across both conformations. The RMSD stabilization, especially the consistent 3-4 Å ligand RMSD, underscores Ang 1-7’s role in mitigating conformational drift, aligning with its protective function in RAS by curbing excessive variability (59). The inactive state experienced a late ∼7 Å peak, suppressed by Ang 1-7, suggests a preventive mechanism against structural instability (51), while the active state exhibited partial stabilization at ∼5 Å, despite a 6-7 Å peak, reflects a less rigid control compared to full agonists (60,61). The RMSF shifts, reduced flexibility in ECL1, ECL3, and TM6, and increased flexibility in ICL2, TM7, and TM5/ICL3, indicate a tailored stabilization strategy, potentially enhancing intracellular signaling (62). This contrasts with the uniform rigidity of classical agonists, supporting Ang 1-7’s partial agonistic profile, as suggested by selective pathway activation in literature (63). The dynamic insights from long-timescale MD simulations highlight adaptability of MasR, offering a foundation for its role in cardiovascular protection (6).

### 3.3. Activation Dynamics and Allosteric Networks of MasR Driven by Ang 1-7

The activation mechanism of MasR was assessed via TM distance analyses (Fig4), dihedral angle shifts (Fig5, FigS6), and dynamic residue interaction network (DRIN) metrics (Fig6, FigS7, TableS3). In the inactive state, TM3-TM6 center-of-mass (COM) distance remained stable at 10 Å for the apo form, while Ang 1-7 binding induced a peak at 15 Å, gradually converging toward the apo level after 600 ns (Fig4a). Similarly, TM3-TM7 distance was stable at 20 Å for apo, with Ang 1-7 binding causing a peak at 25 Å, also approaching apo after 600 ns (Fig4a). This suggests that Ang 1-7 triggers transient widening in the inactive conformation without sustaining outward movement. In the active state, TM3-TM6 distance for apo started at 13-14 Å for the first 200 ns, decreased to ∼12 Å between 200-400 ns, and stabilized at 10 Å from 400-1000 ns, indicating a compacting trend. Ang 1-7 binding, however, began at ∼10 Å, dropped to 7-8 Å between 200-400 ns, and returned to 10 Å from 400-1000 ns, reflecting an initial inward shift followed by stabilization (Fig4b). TM5-TM7 distance remained stable at 22-23 Å for apo, while Ang 1-7 binding peaked at 32 Å between 200-400 ns, then stabilized at 27-28 Å; TM3-TM5 distance for apo was stable at 10 Å for the first 500 ns, rising to 12 Å thereafter, whereas Ang 1-7 binding held at 10 Å for the first 200 ns, peaked at 17-18 Å between 200-800 ns, and converged to 12 Å (Fig4b). Additionally, TM5-TM6 and TM6-TM7 distances increased by 4-5 Å with Ang 1-7 binding throughout the simulation, underscoring a significant TM6 outward displacement. These findings indicate that Ang 1-7 induces transient conformational changes in the inactive state, as evidenced by the 15 Å and 25 Å peaks in TM3-TM6 and TM3-TM7 distances, which revert to apo levels after 600 ns, suggesting a protective role by maintaining stability without triggering activation (64). In the active state, the initial 7-8 Å drop in TM3-TM6 distance with Ang 1-7, followed by a return to 10 Å, contrasts with the apo’s compacting trend, while the 32 Å peak in TM5-TM7 and 17-18 Å peak in TM3-TM5 highlight dynamic widening. The consistent 4-5 Å increase in TM5-TM6 and TM6-TM7 distances with Ang 1-7 binding supports TM6 outward movement, a hallmark of GPCR activation that facilitates G-protein coupling (65). This partial displacement aligns with the role of Ang 1-7 as a partial agonist, modulating rather than fully driving the activation process (66).

**Figure 4.**
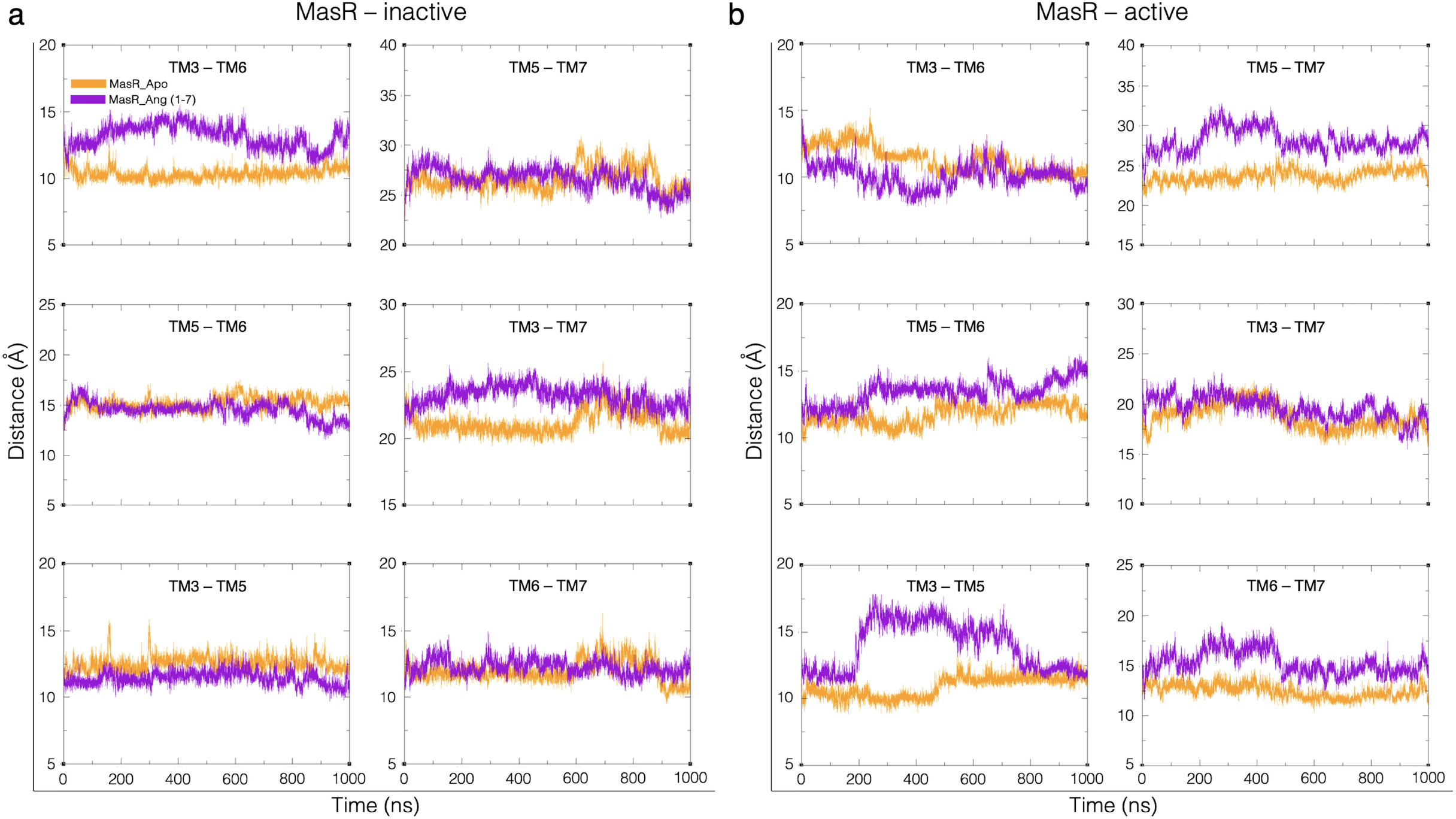
Center of mass (COM) distance analysis of intracellular transmembrane helices of MasR during 1000 ns MD simulations. a) Inactive conformation, (b) Active conformation. Orange lines represent the apo form; purple lines indicate Ang 1–7-bound MasR. COM distances were calculated between key intracellular helices (e.g., TM3–TM6, TM5–TM6, TM6–TM7).

**Figure 5.**
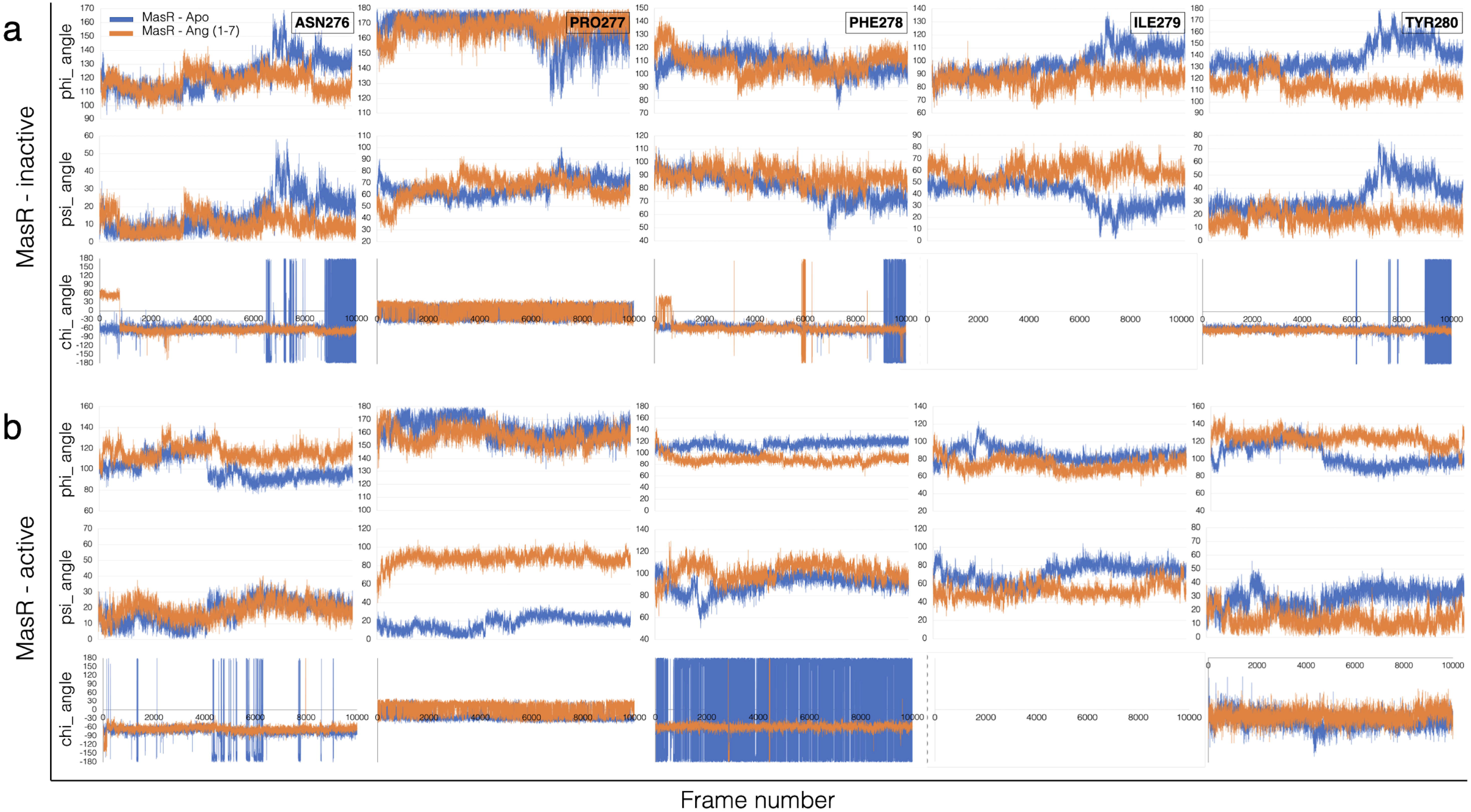
Dihedral angle dynamics of the NPxxY micro-switch motif in MasR during 1000 ns simulations. a) Inactive conformation, b) Active conformation. Time-dependent changes in backbone φ and ψ, and side-chain χ dihedral angles of the NPxxY motif are plotted for apo (blue) and Ang 1–7-bound (orange) conditions.

The NPxxY micro-switch exhibited distinct dihedral dynamics under Ang 1-7 influence, shedding light on its role in MasR activation. In the inactive state, ASN276^7.49^ displayed pronounced φ and ψ angle fluctuations in the apo form, which Ang 1-7 binding reduced, with no significant change until 600 ns, after which φ narrowed by ∼25-20° and ψ by ∼20°. TYR280^7.53^ followed a similar pattern, with φ fluctuations decreasing by ∼20° (0-600 ns) and an additional ∼60° after 600 ns, and ψ reducing by ∼45-50° post-600 ns. ILE279^7.52^ showed a ∼30° φ reduction and a ∼35-40° ψ increase after 600 ns, reflecting dynamic adjustments possibly linked to TM6/H8 regions, while PHE278^7.51^’s ψ increased by ∼20° after 600 ns, hinting at localized shifts. The absence of significant χ1 angle differences (>20°) across all residues underscores that side-chain rotations remain largely unaffected by Ang 1-7 in this state (Fig5a). This late-stage stabilization suggests that Ang 1-7 constrains excessive conformational variability, potentially maintaining the inactive state’s protective configuration (59). This stabilization is mirrored by the NPxxY RMSD, which remained stable at 0.5-0.6 Å with Ang 1-7 binding throughout the simulation, while apo RMSD escalated to 2.5 Å after 600 ns from an initial 0.5-1 Å, highlighting Ang 1-7’s role in maintaining structural order without triggering outward movement (FigS6). In the active state, the NPxxY dynamics shifted toward greater complexity. ASN276^7.49^’s φ angles, initially less variable in apo, increased by ∼20° after 400 ns with Ang 1-7, suggesting enhanced flexibility that may facilitate conformational transitions. TYR280^7.53^’s φ rose by ∼40° after 500 ns, while ψ decreased by ∼25-30° over 0-1000 ns, indicating a balance between flexibility and stabilization that supports partial activation. PRO277^7.50^ exhibited a striking ∼70° ψ increase throughout 0-1000 ns, a significant shift likely driving TM6 outward movement, while PHE278^7.51^’s φ decreased by ∼20° over the same period, contributing to localized stability. ILE279^7.52^’s ψ decreased by ∼20° over 0-1000 ns, further stabilizing the motif. Again, χ_1_ angles showed no substantial differences (>20°), confirming minimal side-chain impact (Fig5b). The NPxxY RMSD remained stable at 0.5 Å with Ang 1-7 binding, while apo maintained 1.5 Å throughout 0-1000 ns (FigS6b), reflecting stabilization amid dynamic changes consistent with partial agonism (67).

DRIN analysis (Fig6, FigS7, TableS3) further elucidated the impact of Ang 1-7 binding on allosteric communication in MasR, highlighting a ligand-induced reconfiguration of key signaling hubs. In the inactive state, betweenness centrality (ΔBC) increased significantly (>±3σ) for critical residues, including TYR280^7.53^ and PHE278^7.51^ within the NPxxY motif, aligning with their late-stage dihedral stabilization (φ and ψ narrowing post-600 ns, Fig5a) and RMSD stability below 0.6 Å (FigS6a), suggesting a stabilizing role (Fig6a, TableS1). Shortest path length (ΔL) increases were noted, indicating altered communication efficiency. In the active state, ΔBC increased notably (>±3σ) for TM5 residue VAL198^5.47^ and H8 residue LEU295^8.58^, with additional >±2σ enhancements for TM6-associated PRO243^6.50^, LEU247^6.54^ and TYR248^6.55^ reflecting improved communication efficiency within these helices and correlating with the observed TM5-TM6 and TM6-TM7 distance expansions (Fig4b, Fig6b, TableS1). ΔL decreases, particularly around TM7 residue ILE279^7.52^, further supported this enhanced connectivity. Snake diagrams (FigS7) visualized these centrality shifts, identifying TYR280^7.53^, PHE278^7.51^, and PHE237^6.44^ as prominent *hot spots* in the ligand-bound state, reinforcing their roles in conformational transmission. Importantly, MasR lacks canonical micro-switch motifs (CWxP, DRY, PIF), yet DRIN results suggest the potential presence of non-canonical equivalents. For instance, the region PHE240^6.47^– PRO243^6.50^ on TM6, showing >±2σ ΔBC shifts (notably PRO243^6.50^), may mimic CWxP’s pivot-like function. ILE126^3.46^–VAL128^3.48^, with >±2σ ΔBC in the active state, could serve as a DRY-like scaffold, while PRO201^5.50^–PHE237^6.44^, spanning TM5–TM6 with >±2σ ΔBC gains, might act as a PIF-like relay, though further validation is needed to confirm these roles based on DRIN data.

**Figure 6.**
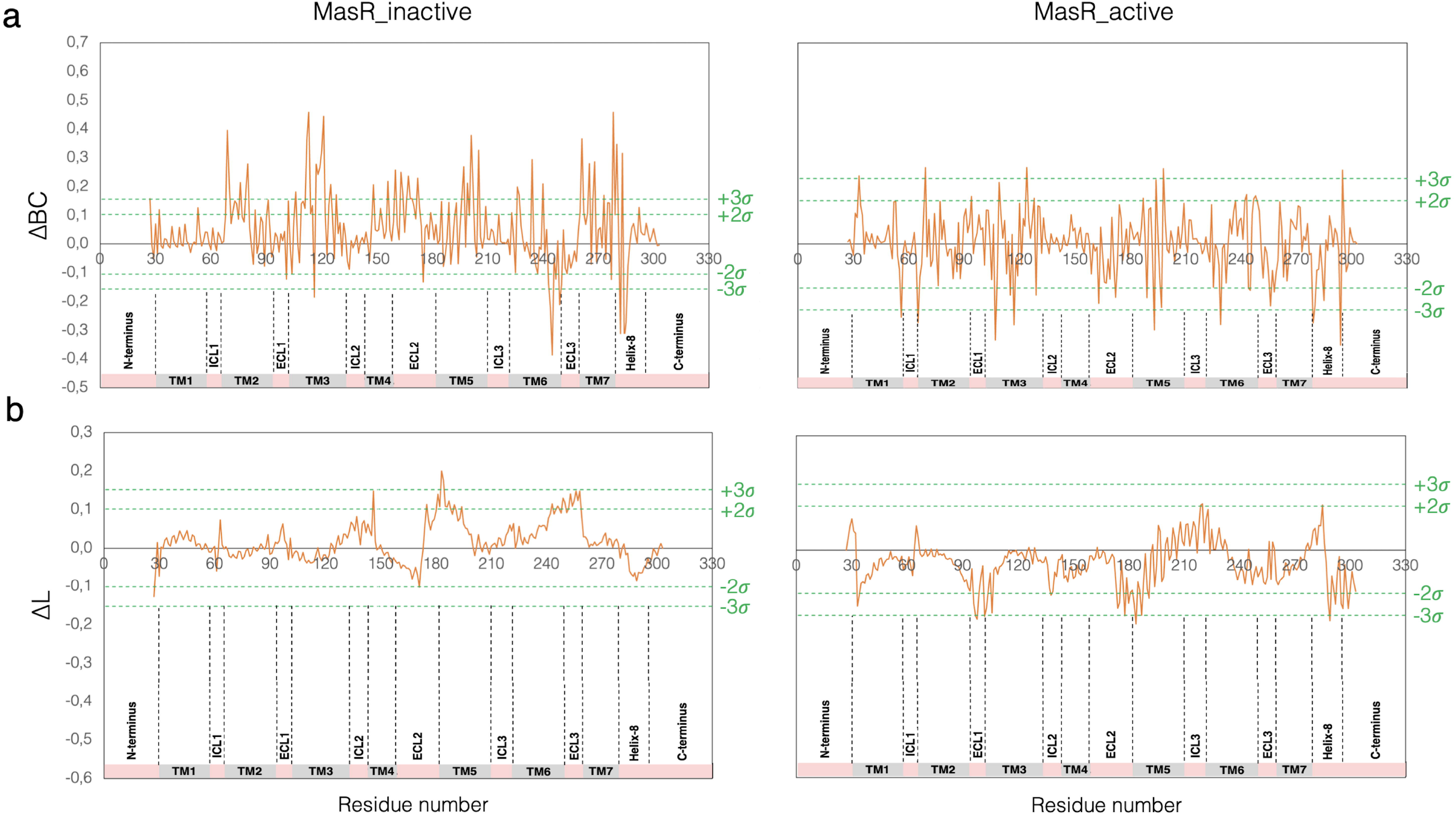
Network-based analysis of MasR reveals differential centrality and communication efficiency induced by Ang 1–7 binding. a) ΔBetweenness centrality (ΔBC) plots for inactive (left) and active (right) MasR. Values represent the difference in residue-level BC between Ang 1–7-bound and apo forms, averaged over simulation replicates and presented with standard deviations. b) ΔShortest path length (ΔL) for inactive (left) and active (right) MasR, showing how ligand binding alters the communication efficiency between residues.

The integrated findings from TM distance analyses, NPxxY dihedral dynamics, and DRIN metrics reveal a multifaceted activation mechanism in MasR modulated by Ang 1-7. In the inactive state, the transient widening of TM3-TM6 and TM3-TM7 distances (Fig4a), coupled with late-stage dihedral stabilization of NPxxY residues (e.g., TYR280^7.53^, PHE278^7.51^) after 600 ns (Fig5a) and RMSD stability at 0.5-0.6 Å (FigS6a), suggests that Ang 1-7 constrains conformational variability, aligning with its protective role in RAS by preventing structural drift (48,59). The increased ΔBC for TYR280^7.53^ and PHE278^7.51^ (Fig6a) supports this stabilizing effect, consistent with anti-inflammatory and vasodilatory functions (68). In the active state, the 4-5 Å TM6 outward displacement (Fig4b), driven by the ∼70° ψ increase in PRO277^7.50^ (Fig5b) and RMSD stability at 0.5 Å (FigS6b), is complemented by elevated ΔBC in TM5 (VAL198^5.47^), TM6 (PRO243^6.50^, LEU247^6.54^, TYR248^6.55^), and H8 (LEU295^8.58^) (Fig6b), alongside ΔL reductions around ILE279^7.52^. This indicates a partial activation that enhances signaling efficiency, yet lacks the full conformational commitment seen in canonical GPCRs [34, 39]. The absence of χ1 angle changes (>20°) (Fig5) and the selective nature of these shifts align with Ang 1-7’s partial agonism, which selectively activates pathways (e.g., ERK1/2, PI3K/Akt) without fully engaging classical G protein signaling (69). The potential non-canonical micro-switch candidates (PHE240^6.47^–PRO243^6.50^, ILE126^3.46^–VAL128^3.48^, PRO201^5.50^–PHE237^6.44^) identified via DRIN (>±2σ ΔBC) further suggest novel regulatory mechanisms in this orphan-type GPCR, though validation is pending (22).

### 3.4. Energetic and Network Connectivity Profiles of MasR Modulated by Ang 1-7

The energetic landscape of Ang 1-7 binding to MasR was characterized using gmx_MMPBSA analysis (Fig7, TableS4), while network connectivity was explored through communication propensity (CP, FigS3), dynamic cross-correlation (DCC, FigS4), and principal component analysis (PCA, FigS5). In the inactive state, the total binding free energy (ΔGbind) was unfavorable at 25.62 kcal/mol, driven by a positive ΔEEL (39.74 kcal/mol) and ΔGSOLV (38.52 kcal/mol), partially offset by ΔVDWAALS (−52.64 kcal/mol) (Fig7a, TableS4). In the active state, ΔGbind became favorable at -13.99 kcal/mol, with enhanced ΔVDWAALS (−55.04 kcal/mol) and reduced ΔEEL (2.90 kcal/mol) contributions (Fig7b, TableS4), indicating a thermodynamically preferred binding mode (70).

**Figure 7.**
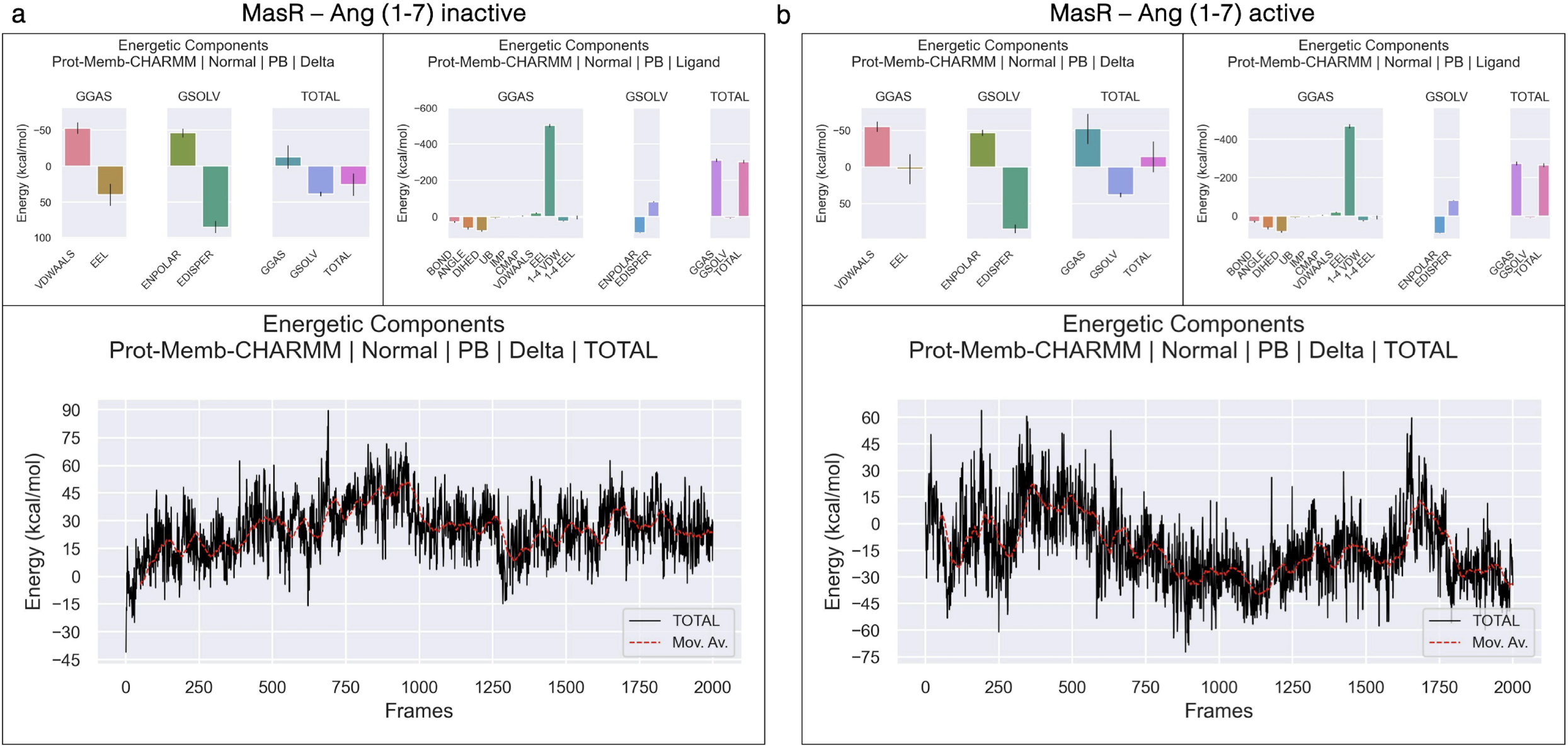
Binding free energy decomposition of Ang 1–7 interacting with inactive and active MasR conformations based on gmx_MMPBSA analysis. a) Energy components for the inactive-state MasR; b) corresponding components for the active-state MasR. Bar plots display individual contributions of van der Waals interactions (VDWAALS), electrostatics (EEL), polar solvation energy (ENPOLAR), nonpolar solvation energy (EDISPER), and the total binding free energy (TOTAL). Line plots show the frame-wise variation of TOTAL energy over the simulation trajectory, with red dashed lines representing the moving average.

The integrated PCA (FigS5), DCC (FigS4), and CP (FigS3) analyses provide a comprehensive view of how Ang 1-7 binding reshapes MasR network connectivity. The PCA results, with PC2 increasing from 9% to 19% in the active state with Ang 1-7, indicate an expanded conformational landscape, suggesting that the ligand induces diverse substates that enhance functional plasticity. This is corroborated by DCC findings, where the transition from strong positive correlations (Cij > 0.7) in Apo to heterogeneous correlations (Cij ≈ 0.3-0.5) with increased anti-correlations (−1.0 to 1.0) in the presence of Ang 1-7, particularly in the active state (Cij ≈ 0.5-0.75), reflects a shift to a more heterogeneous dynamic network. The CP data further support this, with Ang 1-7 elevating communication propensity from below 5 Å^2^ to 5-10 Å^2^, indicating a selective enhancement of inter-residue interactions that modulates rather than fully activates the receptor.

These changes align with literature on GPCRs, where partial agonists induce intermediate conformational states to fine-tune signaling (71). The increased PC2 variance, DCC heterogeneity (Cij ≈ 0.3-0.5 to 0.5-0.75 with anti-correlations), and CP increase (5-10 Å^2^) with Ang (1-7) suggest an allosteric modulation that disrupts coordinated motions, consistent with molecular dynamics studies showing ligand-induced dynamic shifts in GPCRs (72). The CP range of 5-10 Å^2^ and the regulatory pattern of connectivity may be linked to cardiovascular protective effects, as modulated receptor activity could mitigate overactivation (55). The Apo system’s minimal connectivity (below 5 Å^2^) and disorganized active-state dynamics underscore the necessity of ligand binding for network integrity. This ligand-dependent plasticity suggests that Ang 1-7 shapes the signaling of MasR through enhanced flexibility and selective connectivity, differing from full agonists (73). However, these insights are based on simulation data, and experimental validation (e.g., NMR or crystallography) and residue-specific studies are needed to confirm the molecular mechanisms.

Ang 1-7 binding to MasR increases conformational diversity and selectively enhances inter-residue communication propensity, as evidenced by PCA, DCC and CP analyses. This modulatory effect stabilizes and diversifies MasR dynamics, suggesting a protective regulatory role within the RAS (6).

### 3.5. Integrated Mechanistic Insights into Ang 1-7 Modulation of MasR

Our integrative analyses construct a comprehensive mechanistic framework for Ang 1-7’s modulation of MasR, drawing on ligand binding (Fig1-2, TableS2-S3), structural dynamics (Fig3, FigS1-S2), activation mechanisms (Fig4-5-6, FigS6-S7, TableS1), and energetic profiles (Fig7, TableS4, FigS3-S4-S5). In the inactive state, Ang 1-7 stabilizes MasR with moderate H-bond (35.45%), salt bridge (6%), and hydrophobic (53%) occupancies, and an unfavorable ΔGbind (25.62 kcal/mol), suggesting a baseline protective role (6,48,59). In the active state, enhanced occupancies (52.23%, 10%, 84%) and a favorable ΔGbind (−13.99 kcal/mol) reflect increased stability, distinct conformations at ICL2 and TM6 (Fig3b, d) and NPxxY stabilization (Fig5, FigS6) (74,75).

The reduced conformational heterogeneity (FigS2) and network coordination (Fig6, FigS3-S4-S5) in the active state support the partial agonism of Ang 1-7, contrasting with the full activation (63,67,71). Orphan-type GPCR nature of MasR, lacking PIF, E/DRY, or CWxP motifs (22), and its reliance on NPxxY, suggest a unique activation pathway, potentially expandable with novel non-canonical micro-switch candidates (e.g., PHE240^6.47^–PRO243^6.50^, ILE126^3.46^–VAL128^3.48^, PRO201^5.50^–PHE237^6.44^ residues) pending further DRIN/PCA analysis (76). This partial activation aligns with the role of MasR in vasodilation and anti-inflammatory responses within RAS (8,59), offering therapeutic potential for cardiovascular diseases. Unlike short-timescale studies (23), our 1-µs simulations capture these dynamic effects, revealing the plasticity of MasR. The stabilization in active TM6 and H8 regions (65) could guide targeted agonists to enhance protective signaling, addressing gaps in RAS modulation (77). Future experimental validation, such as mutagenesis of key residues (e.g., VAL128^3.48^, PHE237^6.44^, PRO243^6.50^), could refine this framework.

## 4 CONCLUSION

This study provides a comprehensive mechanistic framework for understanding how Ang 1-7 modulates the structure and activation of the MasR, an orphan-type GPCR. Using long-timescale (1 µs) MD simulations of AlphaFold-predicted active and inactive MasR conformations, we demonstrate that Ang 1-7 binding induces distinct state-dependent effects: it stabilizes the inactive conformation by constraining NPxxY dynamics and suppressing late-stage conformational fluctuations, while partially activating the receptor in the active state via TM6 displacement, dynamic micro-switch reorganization, and enhanced network connectivity.

Energetic analyses (MM/PBSA) revealed a thermodynamically favorable binding mode in the active state, supported by increased occupancy of hydrogen bonds, salt bridges, and hydrophobic contacts. DRIN metrics and PCA further indicated a transition toward a more connected and flexible network architecture, consistent with partial agonism. Notably, in the absence of classical micro-switch motifs (CWxP, DRY, PIF), our results suggest non-canonical regulatory elements (e.g., PHE240^6.47^–PRO243^6.50^, ILE126^3.46^–VAL128^3.48^, PRO201^5.50^–PHE237^6.44^) that may compensate structurally and dynamically for MasR activation.

These findings expand the molecular understanding of MasR activation and its modulation by Ang 1-7, supporting its classification as a partially activated receptor with therapeutic potential in cardiovascular and renal pathophysiology. This work also lays the foundation for future experimental studies, including mutagenesis and structure-based drug design efforts targeting MasR-specific regulatory hotspots.

## DATA AND CODE AVAILABILITY

The codes used for analyzing micro-switch dihedral angles and hydrophobic interactions during molecular dynamics simulations are publicly available on GitHub at the following links:

https://github.com/eygpcr/microswitches_dihedral_angles https://github.com/eygpcr/hydrophobic_interactions

Trajectory files of the simulations and related data for MasR can be accessed through the link https://doi.org/10.5281/zenodo.15781385.

## SUPPORTING MATERIAL

Supporting Materials and Methods, which illustrate detailed analyses, including ligand-receptor interactions, structural dynamics and activation mechanisms, include 7 figures, 4 tables and a data file for the movies (https://doi.org/10.5281/zenodo.15781385).

## AUTHOR CONTRIBUTIONS

E.Y. designed and performed the research and wrote the manuscript. S.D., E.E., and N.Y. designed the research and wrote the manuscript.

ACKNOWLEDGEMENTS

This research was conducted as part of Dr. Ekrem Yasar’s PhD dissertation at Akdeniz University Medical School, under the supervision of Prof Nazmi Yaras. This work was supported by Akdeniz University Scientific Research Projects Found [Grant No. TDK-5449, 2021].

## Declaration of generative AI and AI-assisted technologies in the writing process

During the preparation of this work the author(s) used *ChatGPT* [OpenAI, L.L.C., San Francisco, CA] as well as *DeepL Write* [DeepL SE, Cologne, Germany] in order to writing process for spelling and grammar corrections. After using this tool/service, the author(s) reviewed and edited the content as needed and take(s) full responsibility for the content of the published article.

