## Supplemental Information for "Angiotensin 1-7 Modulates the Dynamics and Activation of the Proto-oncogene Mas Receptor"

1. **SUPPLEMENTARY METHOD**
   1.1. **Communication Propensity (CP)**

Dynamic communication between different regions of a protein is critical for maintaining structural stability, regulating allosteric signaling, and enabling functional activation, particularly in complex systems such as G protein-coupled receptors (GPCRs). Communication Propensity (CP) is a quantitative metric that assesses the stability of spatial relationships between residue pairs by measuring fluctuations in their inter-residue distances over the course of molecular dynamics (MD) simulations. This approach is grounded in the concept of signal transmission pathways, wherein chains of dynamically interacting residues relay information across different regions of a protein. CP quantifies these fluctuations: residue pairs that exhibit greater variability in their Cα–Cα distances are considered less stable and less efficient in communication, whereas pairs with smaller fluctuations are indicative of more consistent and efficient signal transmission. The extent of these distance fluctuations directly influences the efficiency of communication, with higher variability associated with reduced signal propagation speed. Mathematically, CP between residues i and j is defined as:

$\boldsymbol{CP}\mathbf{=}\left\langle\left( \boldsymbol{d}_{\boldsymbol{ij}}\boldsymbol{-}\boldsymbol{d}_{\boldsymbol{ij,ave}} \right)^{\mathbf{2}} \right\rangle$	(1)

where $d_{ij}$is the instantaneous distance between the Cα atoms of residues i and j at each frame, and $d_{ij,ave}$represents the time-averaged distance over the trajectory. CP values range from 0 to 1, with higher values indicating a greater propensity for efficient communication and a stronger involvement in dynamic signaling pathways (Chennubhotla & Bahar, 2007; Morra et al., 2009; Penkler & Tastan Bishop, 2019). Lower CP values correspond to weaker communication, characterized by larger fluctuations in inter-residue distances.

The fundamental premise underlying CP analysis is that regions maintaining stable relative positions during protein dynamics are more likely to participate in efficient allosteric signal transmission or structural rearrangements required for functional activity. Thus, CP provides a powerful means of identifying dynamically coupled networks within proteins that may underlie critical functional transitions, such as ligand-induced activation of receptors.

In this study, CP analysis was utilized to evaluate the signal transmission efficiency between residue pairs in MasR with and without ligand binding (Ang 1-7) conditions. CP values were computed based on the squared fluctuations of Cα–Cα distances across 2,000 frames of each MD trajectory, resulting in unnormalized values typically ranging from 0 to 25 Å². This distribution reflects the extent of dynamic stability between residue pairs, with lower CP values indicating stronger communication pathways and higher values corresponding to more flexible, unstable connections. CP was computed using MDM-TASK (Amamuddy et al., 2021; Brown et al., 2017) identifying residue pairs with stable interactions that enhance communication pathways modulated by ligand binding, thus revealing the dynamic signaling architecture of MasR.

1.2. **Dynamic Cross Correlation (DCC)**

Dynamic cross correlation (DCC) analysis is a widely utilized method for characterizing the coordinated or opposing motions between different regions of a protein structure during molecular dynamics (MD) simulations. DCC provides quantitative insights into how positional fluctuations of atoms or residues are synchronized over time, offering a detailed map of dynamic couplings within the protein. This information is crucial for understanding the structural and functional interdependence of different regions, particularly in dynamic systems such as G protein-coupled receptors (GPCRs).

DCC measures the cross-correlation between the fluctuations of two residues, assessing whether their movements are positively correlated (moving in the same direction), negatively correlated (moving in opposite directions), or uncorrelated. Mathematically, the cross-correlation coefficient *C_ij_* between residues *i* and *j* is calculated as:

$\boldsymbol{C}_{\boldsymbol{i,j}}\boldsymbol{=}\frac{\left\langle\left( \boldsymbol{r}_{\boldsymbol{i}}\boldsymbol{-}\left\langle\boldsymbol{r}_{\boldsymbol{i}} \right\rangle\right)\boldsymbol{\cdot}\left( \boldsymbol{r}_{\boldsymbol{j}}\boldsymbol{-}\left\langle\boldsymbol{r}_{\boldsymbol{j}} \right\rangle\right) \right\rangle}{\sqrt{\left( \left\langle\boldsymbol{r}_{\boldsymbol{i}}^{\boldsymbol{2}} \right\rangle\boldsymbol{-}\left\langle\boldsymbol{r}_{\boldsymbol{i}} \right\rangle^{\boldsymbol{2}} \right)}\left( \left\langle\boldsymbol{r}_{\boldsymbol{j}}^{\boldsymbol{2}} \right\rangle\boldsymbol{-}\left\langle\boldsymbol{r}_{\boldsymbol{j}} \right\rangle^{\boldsymbol{2}} \right)}$ (2)

where $\boldsymbol{r}_{\boldsymbol{i}}$ and $\boldsymbol{r}_{\boldsymbol{j}}$ are the instantaneous positional vectors of residues *i* and *j* at each frame, $\left\langle\boldsymbol{r}_{\boldsymbol{i}} \right\rangle$ and $\left\langle\boldsymbol{r}_{\boldsymbol{j}} \right\rangle$ denote their respective time-averaged positions over the trajectory, and the bracket-enclosed quantities represent averages computed over all frames (Bowerman & Wereszczynski, 2016; Kasahara et al., 2014). The resulting DCC values range from −1 to +1, where +1 indicates perfectly correlated motion, −1 indicates perfectly anticorrelated motion, and values near zero suggest independent movements.

The fundamental principle of DCC analysis is that regions exhibiting strong correlation or anticorrelation are dynamically coupled and may cooperatively participate in structural transitions, allosteric regulation, or ligand-induced conformational changes. Therefore, DCC analysis serves as a powerful tool for elucidating the functional architecture and dynamic connectivity within proteins.

In this study, DCC was employed to assess the dynamic correlations between residue pairs in MasR across 2,000 frames of each MD trajectory, using MDM-TASK (Amamuddy et al., 2021; Brown et al., 2017) for with and without ligand binding (Ang 1-7) conditions. This analysis identified regions with enhanced or diminished coupling due to ligand interactions, elucidating how ligand binding modulates the structural dynamics and communication networks of MasR.

1.3. **Principal Component Analysis (PCA)**

Principal component analysis (PCA) is a statistical technique commonly employed to simplify and interpret the large and complex datasets generated from molecular dynamics (MD) simulations. PCA reduces the dimensionality of multivariate data by transforming a set of correlated variables, such as atomic positional fluctuations, into a new coordinate system characterized by orthogonal (linearly uncorrelated) axes, known as principal components. Each principal component captures a distinct mode of motion within the protein, ranked according to the variance it explains. In molecular dynamics, PCA is typically applied to the time series of atomic coordinates to identify dominant structural fluctuations and conformational transitions that contribute to the protein’s functional behavior.

The transformation relies on the construction of the covariance matrix C, where each element *C_ij_* is defined as:

$\boldsymbol{C}_{\boldsymbol{i,j}}\boldsymbol{=}\left\langle\left( \boldsymbol{x}_{\boldsymbol{i}}\boldsymbol{-}\left\langle\boldsymbol{x}_{\boldsymbol{i}} \right\rangle\right)\left( \boldsymbol{x}_{\boldsymbol{j}}\boldsymbol{-}\left\langle\boldsymbol{x}_{\boldsymbol{j}} \right\rangle\right) \right\rangle$	(3)

where $\boldsymbol{x}_{\boldsymbol{i}}$ and $\boldsymbol{x}_{\boldsymbol{j}}$ represent the Cartesian coordinates of atoms *i* and *j*, respectively. The angle brackets indicate time-averaged values calculated over all structures sampled throughout the MD trajectory (Amamuddy et al., 2021; David & Jacobs, 2014). Diagonalization of the covariance matrix yields eigenvectors and eigenvalues, where each eigenvector corresponds to a collective mode of motion, and its associated eigenvalue indicates the magnitude of variance along that mode. Principal components with larger eigenvalues reflect dominant motions that account for substantial conformational variability. By projecting trajectory data onto the principal components, PCA provides a quantitative description of essential dynamics, facilitating the identification of structural rearrangements and dynamic landscapes involved in protein function and ligand interactions.

In this study, PCA was performed on MasR trajectories to extract the primary conformational modes influenced by Ang 1-7 binding. The covariance matrix of Cα atomic positions was computed over 2,000 frames for each MD trajectory using MDM-TASK (Brown et al., 2017). PCA was conducted separately for ligand-free and ligand-bound systems, enabling the identification of key structural fluctuations and providing insight into the dynamic responses of MasR to different ligand binding events.

1.4. **Dynamic Residue Interaction Network (DRIN)**

Dynamic Residue Interaction Network (DRIN) analysis leverages graph theory to model a protein as a network of nodes (residues) and edges (interactions) that evolve over the course of molecular dynamics (MD) simulations. In this representation, each residue is mapped to its C_β_ atom (or C_α_ atom for glycine residues), and an edge is established between two residues if their representative atoms remain within a predefined distance cutoff, typically 6.7 Å, for a significant portion of the simulation. This network-based approach captures the dynamic nature of protein interactions, revealing pathways and communication hubs that mediate structural stability and functional transitions.

DRIN analysis employs two key graph-theoretical metrics to characterize the network: the average betweenness centrality (*BC*) and the average shortest path length (*L_i_*). The average BC of a residue quantifies how often that residue appears on the shortest communication paths between all other pairs of residues, thus reflecting its importance as a communication hub. The average *BC* is calculated as:

$\bar{\boldsymbol{BC}_{\boldsymbol{v}}}\boldsymbol{=}\frac{\boldsymbol{1}}{\boldsymbol{m}}\sum_{\boldsymbol{i=1}}^{\boldsymbol{m}} \sum_{\boldsymbol{s,t\epsilon V}} \frac{\boldsymbol{\sigma}_{\boldsymbol{i}}\left( \boldsymbol{s,t}\mathbf{I}\boldsymbol{v} \right)}{\boldsymbol{\sigma}_{\boldsymbol{i}}\left( \boldsymbol{s,t} \right)}$(4)

where $\sigma_{i}\left( s,tIv \right)$ is the number of shortest paths passing through residue *v* at frame *i*, $\sigma_{i}\left( s,t \right)$ is the total number of shortest paths between residues *s* and *t*, and *m* is the total number of frames (Penkler & Tastan Bishop, 2019; Sethi et al., 2009).

The average shortest path length (*L_i_*) represents the average geodesic distance from a given residue to all other residues in the network, providing a measure of how well-connected a residue is. *Li* is defined as:

$\boldsymbol{L}_{\boldsymbol{i}}\boldsymbol{=}\frac{\boldsymbol{1}}{\boldsymbol{N-1}}\sum_{\boldsymbol{j=1}}^{\boldsymbol{N}} \boldsymbol{L}_{\boldsymbol{ij}}$ (5)

where $L_{ij}$ is the shortest path length between residues *i* and *j*, and *N* is the total number of residues. Shortest paths are computed using algorithms such as Dijkstra’s algorithm based on inter-residue distances.

By analyzing these network properties, DRIN provides insights into the dynamic communication architecture of proteins, enabling the identification of residues that act as critical hubs or bridges within dynamic signaling pathways.

In this study, DRIN analysis was conducted on MasR to map dynamic interaction networks in with and without ligand binding (Ang 1-7) conditions. Networks were constructed from 2,000 frames of each MD trajectory using MDM-TASK (Amamuddy et al., 2021; Brown et al., 2017) treating C_β_ atoms (C_α_ for glycine) as nodes and applying a 6.7 Å cutoff to define edges. Average *BC* and *L_i_* values were computed to identify key residues involved in communication networks and to assess ligand-induced alterations in dynamic signaling structure of MasR.

2. **SUPPORTING MATERIAL**

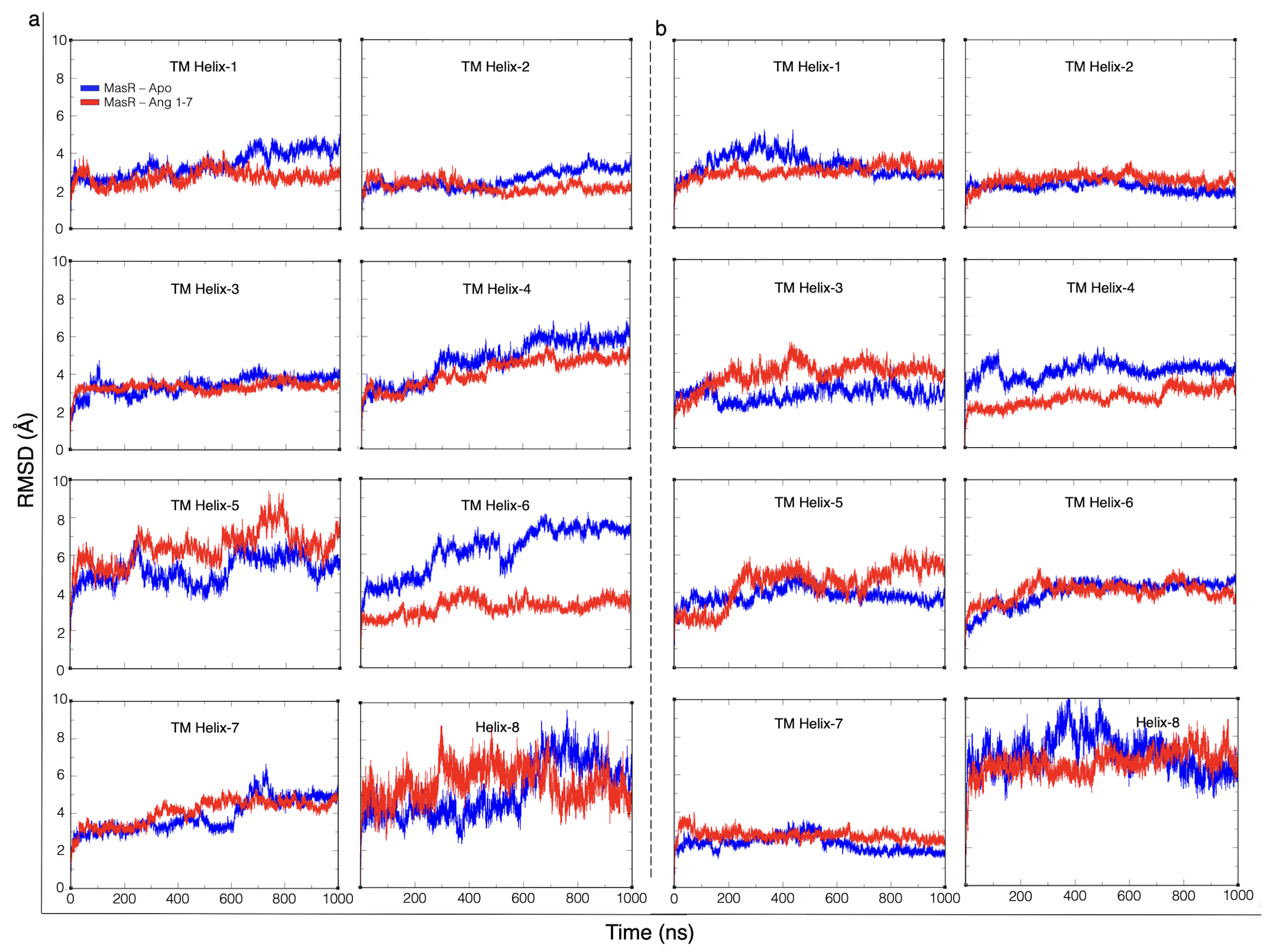

**Figure S1.** **RMSD profiles of individual transmembrane helices (TM1–TM7) and helix 8 (H8) of MasR in inactive and active states during simulations.** a) Inactive-state MasR. RMSD trajectories of TM helices and H8 are shown for apo (blue) and Ang 1–7-bound (red) forms. b) Active-state MasR.

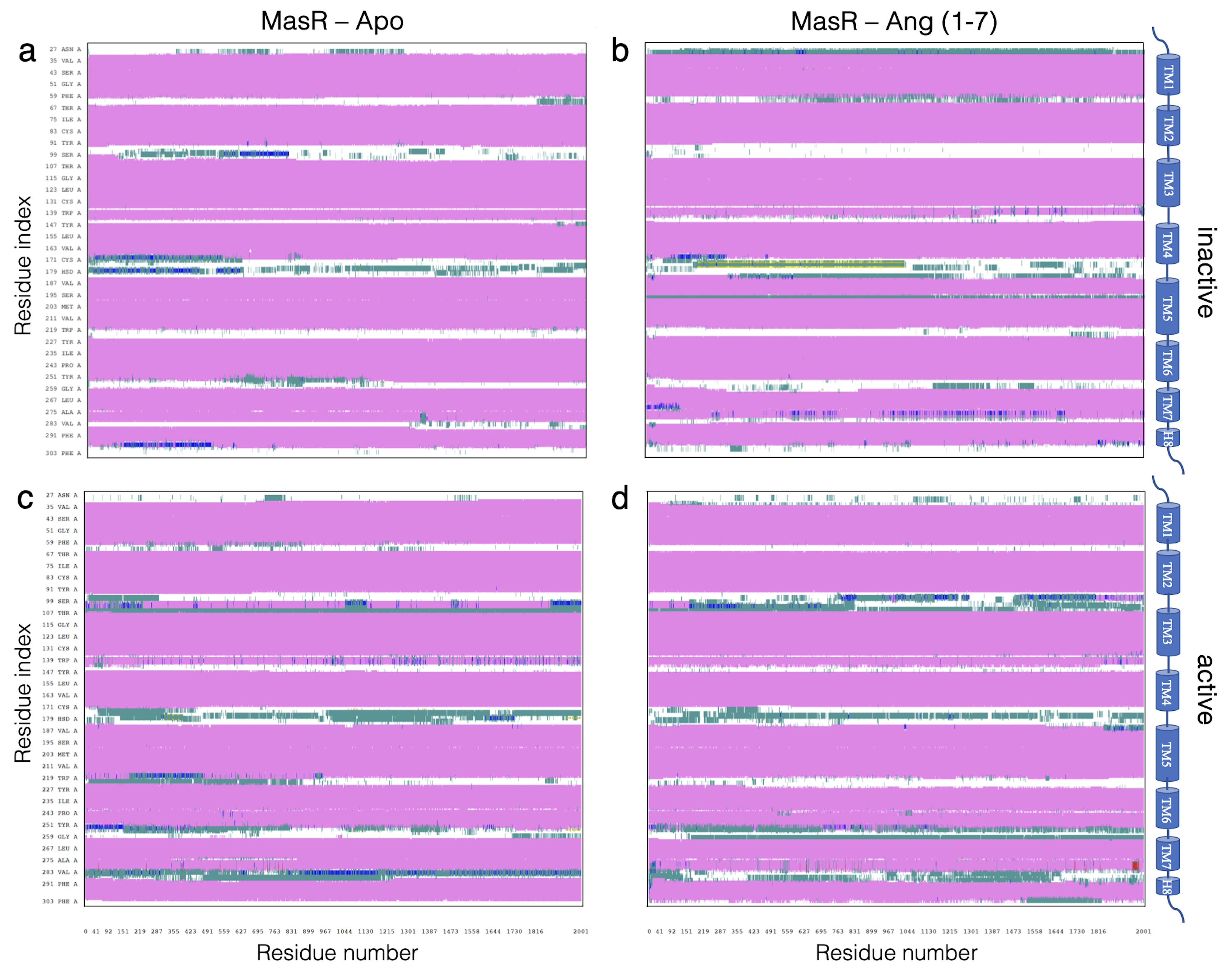

**Figure S2.** **Two-dimensional structural timeline analyses of Mas receptor (MasR) systems during MD simulations in different conformational states.** Residue-wise structural fluctuations over time are visualized using the VMD Timeline plugin. Each row represents a residue, and colors indicate changes in backbone dihedral conformations across time frames. a) Apo-inactive MasR, b) Ang 1–7-bound inactive MasR, c) Apo-active MasR, and d) Ang 1–7-bound active MasR.

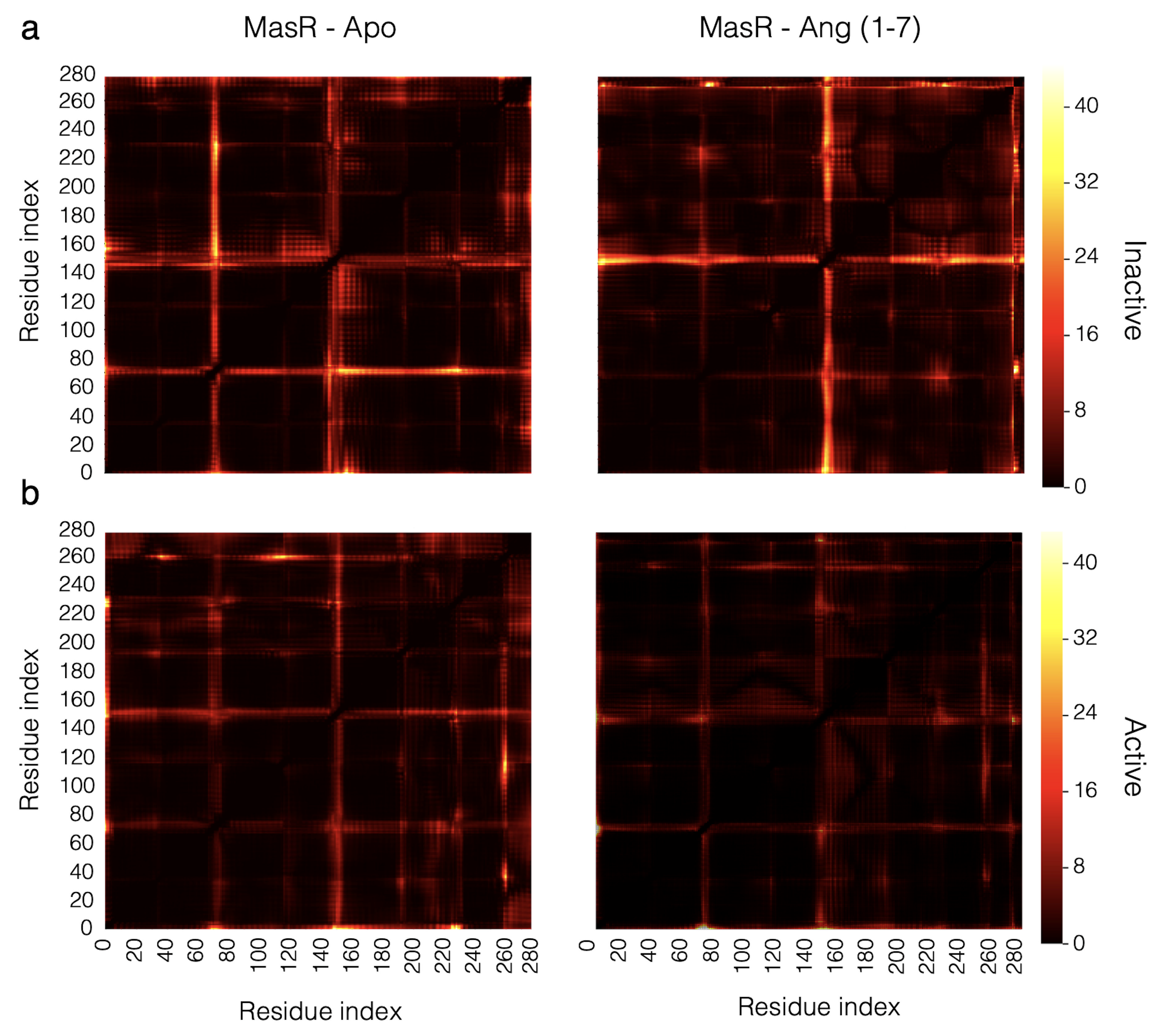

**Figure S3.** **Comparative communication propensity (CP) analysis of MasR in inactive and active states with and without Ang 1–7.** a) CP heatmaps for the inactive-state MasR: apo (left) and Ang 1–7-bound (right). b) CP heatmaps for the active-state MasR: apo (left) and Ang 1–7-bound (right). The color scale indicates the extent of distance fluctuation between residue pairs, with lower values representing more stable communication pathways.

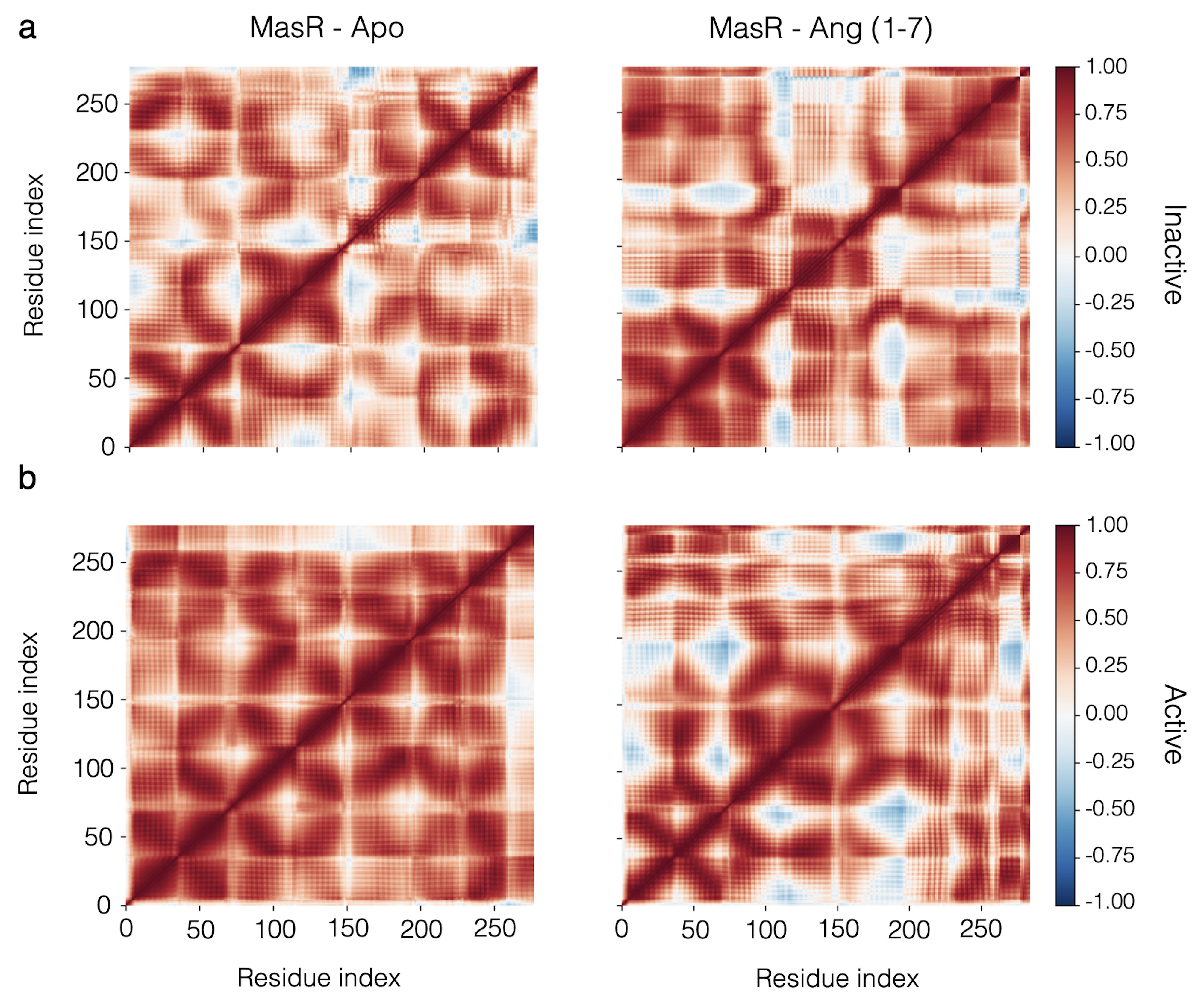

**Figure S4.** **Dynamic Cross-Correlation (DCC) analysis of MasR in inactive and active states with and without Ang 1–7.** Heatmaps represent residue-level correlation patterns during 1000 ns MD simulations. a) Inactive-state apo form (Left), Inactive-state Ang 1–7-bound form (Right), b) Active-state apo form (Left), Active-state Ang 1–7-bound form (Right). The color scale reflects the degree of correlated (red) or anti-correlated (blue) motions between residue pairs during molecular dynamics simulations.

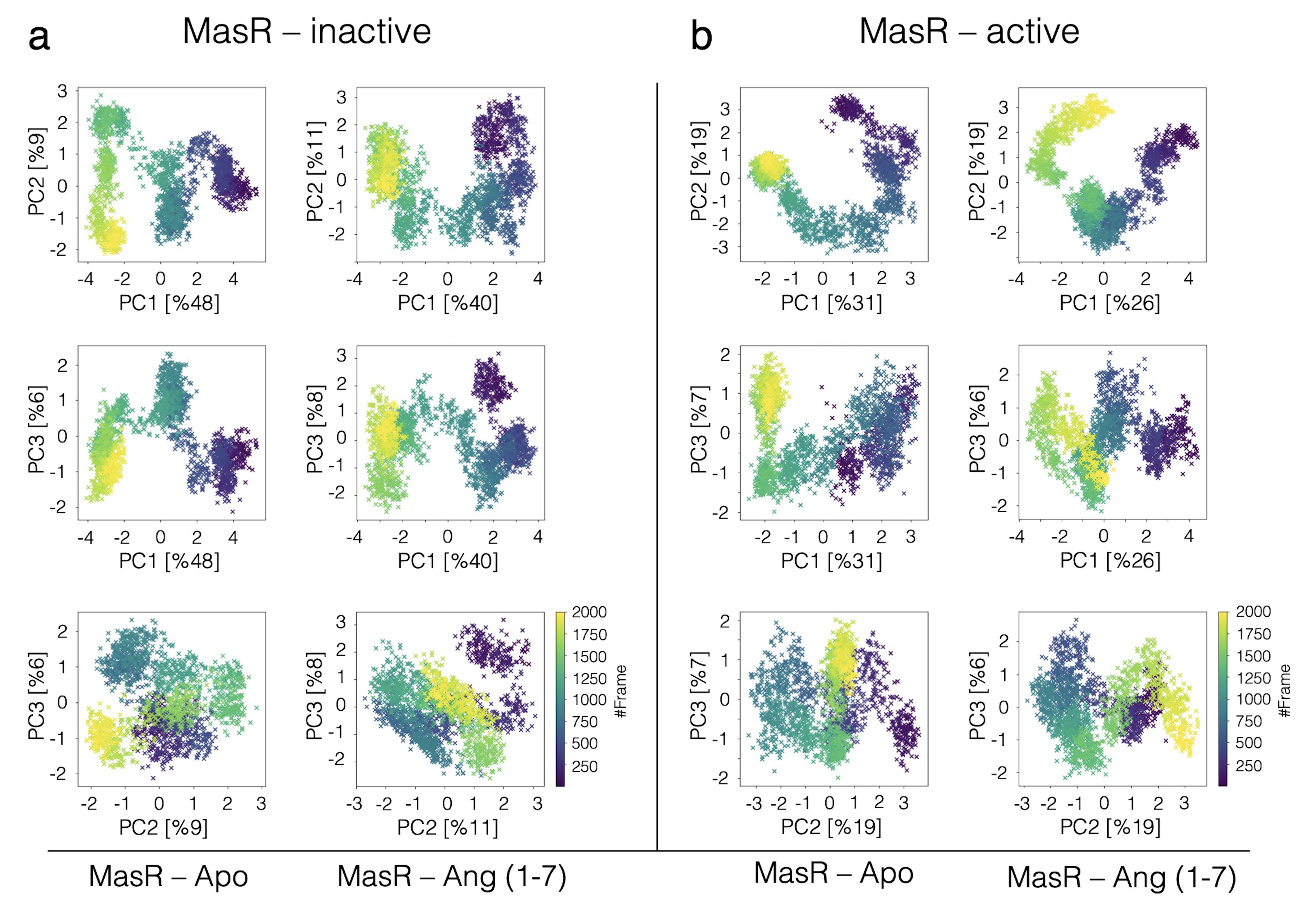

**Figure S5.** **Principal component analysis (PCA) of MasR conformational dynamics in inactive and active states.** a) PCA projections of MD trajectories for the inactive-state MasR in apo and Ang 1–7-bound conditions. b) Equivalent PCA projections for the active-state MasR. Conformational sampling is shown along PC1–PC2, PC1–PC3, and PC2–PC3 axes, with explained variance percentages indicated.

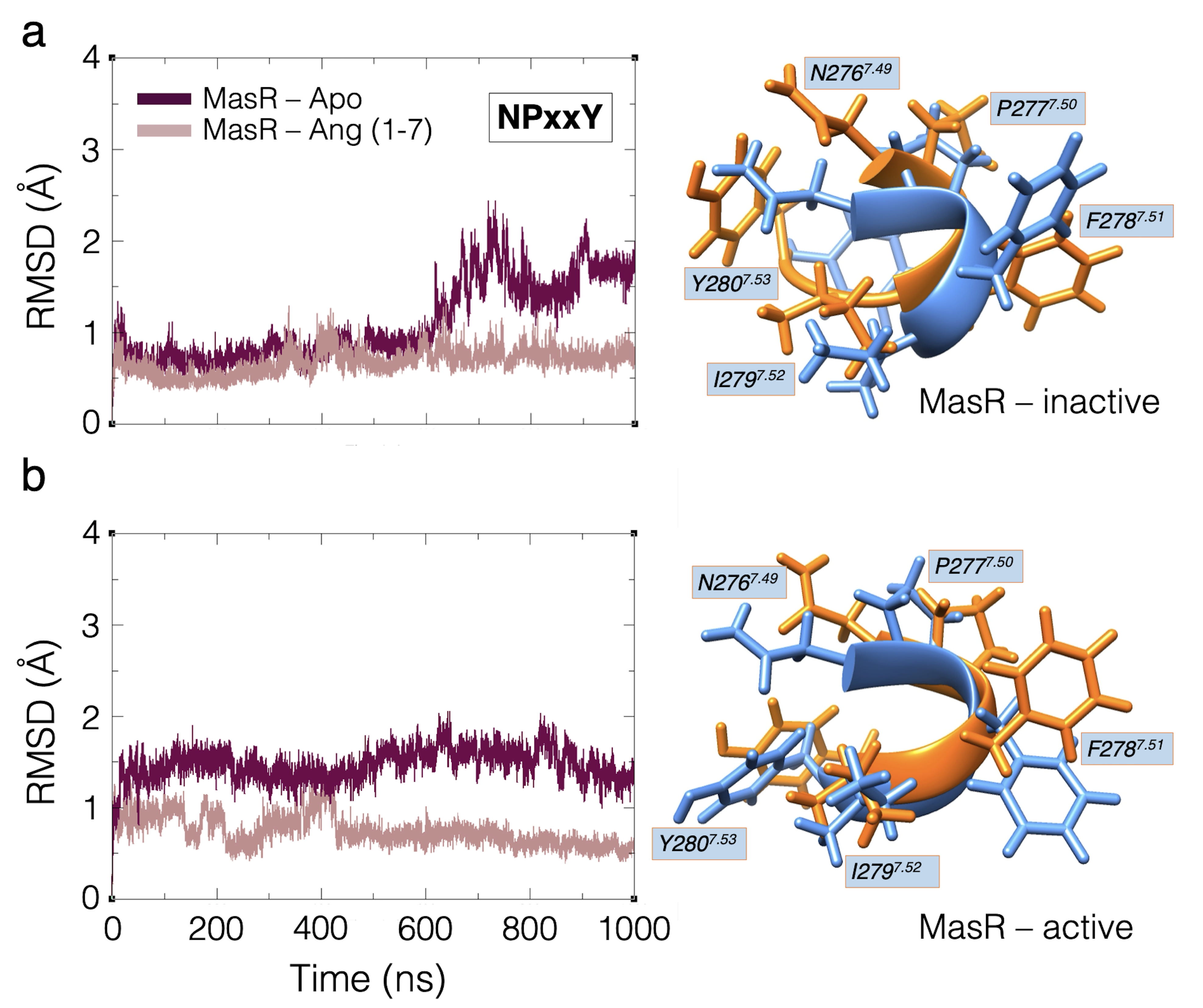

**Figure S6.** **Structural stability of the NPxxY motif in MasR during 1000 ns MD simulations.** a) Inactive conformation, b) Active conformation. Root-mean-square deviation (RMSD) plots of the NPxxY motif are shown for apo (blue) and Ang 1–7-bound (orange) conditions.

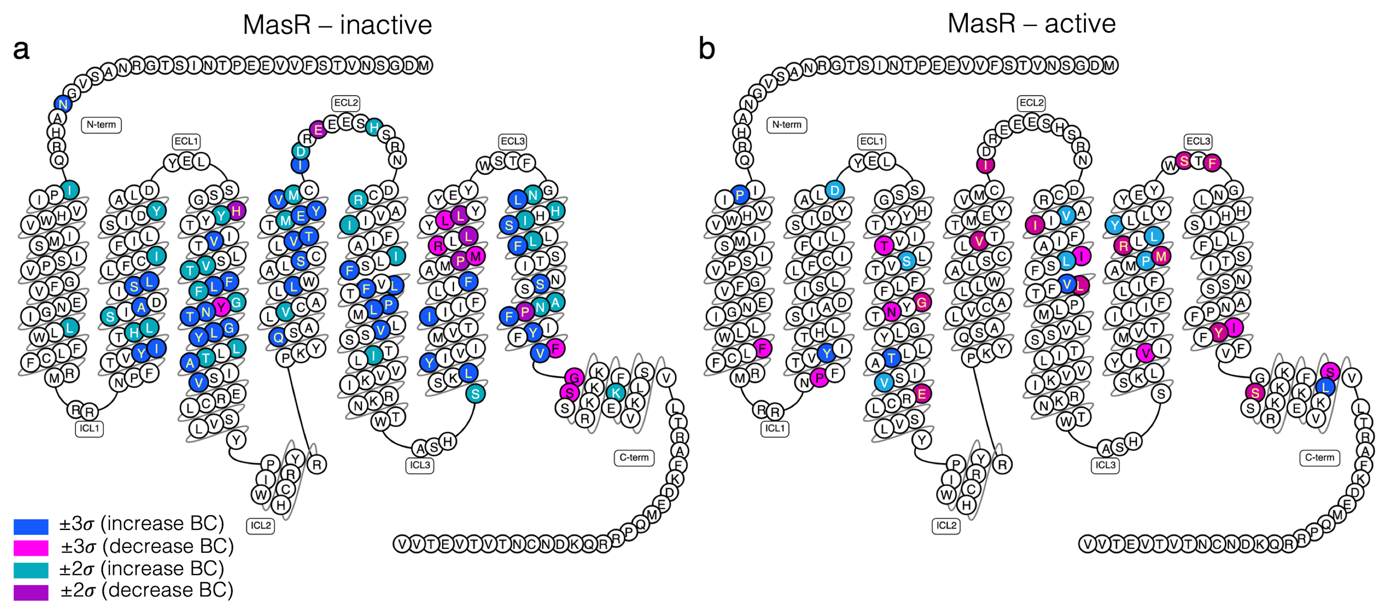

**Figure S7.** **Snake diagram visualization of residue-level network centrality changes in MasR upon Ang 1–7 binding, based exclusively on ΔBetweenness Centrality.** a) Inactive conformation, and b) Active conformation of MasR. Residues are colored according to the change in betweenness centrality (ΔBC) values calculated from dynamic residue interaction network (DRIN) analysis (see Figure 8, TableS3). Residues are color-coded based on changes in BC relative to the mean: blue indicates an increase >3σ, cyan indicates an increase >2σ, magenta indicates a decrease >3σ, and purple indicates a decrease >2σ.

3. **SUPPLEMENTARY REFERENCES**

Amamuddy, O. S., Glenister, M., & Tastan Bishop, Ö. (2021). MDM-TASK-web: A web platform for protein dynamic residue networks and modal analysis. *bioRxiv*, *19*, 5059-5071.

Bowerman, S., & Wereszczynski, J. (2016). Detecting allosteric networks using molecular dynamics simulation. *Methods in enzymology*, *578*, 429-447.

Brown, D. K., Penkler, D. L., Sheik Amamuddy, O., Ross, C., Atilgan, A. R., Atilgan, C., & Tastan Bishop, Ö. (2017). MD-TASK: a software suite for analyzing molecular dynamics trajectories. *Bioinformatics*, *33*(17), 2768-2771.

Chennubhotla, C., & Bahar, I. (2007). Signal propagation in proteins and relation to equilibrium fluctuations. *PLoS computational biology*, *3*(9), e172.

David, C. C., & Jacobs, D. J. (2014). Principal component analysis: a method for determining the essential dynamics of proteins. *Protein dynamics: Methods and protocols*, 193-226.

Kasahara, K., Fukuda, I., & Nakamura, H. (2014). A novel approach of dynamic cross correlation analysis on molecular dynamics simulations and its application to Ets1 dimer–DNA complex. *PLoS One*, *9*(11), e112419.

Morra, G., Verkhivker, G., & Colombo, G. (2009). Modeling signal propagation mechanisms and ligand-based conformational dynamics of the Hsp90 molecular chaperone full-length dimer. *PLoS computational biology*, *5*(3), e1000323.

Penkler, D. L., & Tastan Bishop, Ö. (2019). Modulation of human Hsp90α conformational dynamics by allosteric ligand interaction at the C-terminal domain. *Scientific reports*, *9*(1), 1600.

Sethi, A., Eargle, J., Black, A. A., & Luthey-Schulten, Z. (2009). Dynamical networks in tRNA: protein complexes. *Proceedings of the National Academy of Sciences*, *106*(16), 6620-6625.
